# Emergence of a complex logic gate from the integration of slippage-induced frameshift mechanisms

**DOI:** 10.64898/2026.04.01.715767

**Authors:** Rushik Bhatti, Kayarat Saikrishnan

**Author notes:** Correspondence should be addressed to K.S.

## Abstract

Phase variation is viewed as a simple stochastic ON/OFF switch helping bacteria survive unpredictable environments. In minimal-genome pathogens like *Mycoplasma bovis*, simple sequence repeats (SSRs) introduce frameshifting InDels in key phase-variable genes, such as the Type III restriction-modification *mod* genes, typically assumed to result in binary expression. This study revisits this assumption using a heterologous *E. coli* system and single-cell mEGFP-based fluorescence profiling of *M. bovis mod1* gene fragment containing SSR to determine if "OFF" states are truly silent. We find that the frameshifted construct remains active, showing low-level, heterogeneous expression in a frame-dependent manner, via frameshift suppression. This creates a range of expression rather than a strict binary switch. These findings suggest that phase-variable SSRs can function with upstream switches to form complex XNOR Boolean logic gates. This demonstrates that sophisticated logic gates can emerge directly from coding sequence architecture enhancing diversity and adaptability to promote evolutionary resilience in compact bacterial genomes.

## INTRODUCTION

Phase variation is a powerful bet-hedging strategy used by numerous host-adapted bacteria to evade host defenses and adapt to rapidly fluctuating environments. It is a phenomenon that leads to a high-frequency, stochastic, reversible, and heritable switching of phenotypes (Moxon et al., 1994, 2006; Van Der Woude and Bäumler, 2004). One crucial component driving phase variation is the presence of simple sequence repeats (SSRs). SSRs are known to exhibit a higher propensity for slipped-strand mispairing during DNA replication, leading to high-frequency length polymorphisms or InDel mutations (Levinson and Gutman, 1987a, 1987b). Such phase-variable SSRs (PV SSRs) are extensively found in the coding sequences of host-adapted prokaryotes (De Bolle et al., 2000; Lin and Kussell, 2012). Such PV SSRs are prevalent in host-adapted pathogens with reduced genomes, such as *Mycoplasma* species, where length polymorphisms at PV SSR loci have been implicated in phase variation (Mrázek et al., 2007). *Mycoplasma bovis*, a cell-wall-less bacterium in the class Mollicutes, is a significant bovine pathogen causing chronic respiratory disease, mastititis, arthritis, otitis, and keratoconjunctivitis. Worldwide outbreaks underscore their economic impact on cattle industries, driven by antimicrobial resistance and a lack of effective vaccines (Lysnyansky and Ayling, 2016). With a compact genome of approximately 1 Mbp, it exhibits phase variation and antigenic switching to evade the immune response (Citti et al., 2010; Dybvig and Voelker, 1996; Li et al., 2011; Lysnyansky and Ayling, 2016).

Genera such as *Mycoplasma* and *Ureaplasma* present a unique case in the distribution of restriction-modification (R-M) systems among prokaryotes. Despite their small genomes, this class of bacteria exhibits a high density of R-M systems, ranking second among all analysed phyla in a comparative study of 2,261 prokaryotic genomes (Oliveira et al., 2014). *M. bovis* genomes harbour a rich repertoire of phase-variable Type III R-M systems with mostly dinucleotide (AG)_n_ PV SSR tracts in their *mod* genes. *Mycoplasma* species encode multiple paralogs of phase-variable *mod* genes, often existing as allelic variants with distinct Target Recognition domains (TRD) (Atack et al., 2018). Frameshifting InDel mutations at PV SSR loci in *mod* CDS can shift the reading frame, resulting in reversible functional loss and altered global DNA methylation patterns (Srikhanta et al., 2005). Such stochastic switching of Mod expression can possibly generate phenotypic heterogeneity within the *Mycoplasma* population.

Recent work on frameshifting InDels in protein-coding genes has revealed that the deleterious impact of frameshifts can be substantially bypassed by RNA polymerase slippage or ribosomal slippage, enabling the bypass of frameshift lesions and the production of full-length functional proteins (Rockah-Shmuel et al., 2013). Complementary theoretical and experimental studies further suggest that phenotypic mutations, including ribosomal frameshifts and stop-codon readthrough, generate transient proteoforms that can buffer genetic defects, expand functional diversity, and shape long-term protein evolution (Goldsmith and Tawfik, 2009; Romero Romero et al., 2022). In this context, frameshifted *mod* gene in host bacteria may not be strictly null alleles but instead give rise to low-level or condition-dependent expression of functional methyltransferase variants via frameshift suppression. However, direct experimental tests of such suppression, particularly for phase-variable Type III mod from host-adapted pathogens like *M. bovis*, are still lacking.

Here, we investigate frameshift suppression of a frameshifted Type III *mod1* gene from *Mycoplasma bovis* PG45, expressed in *E. coli*, using mEGFP reporter-based single-cell profiling. By cloning phase-variable *mod* genes containing dinucleotide (AG)_n_ repeat tracts, the study aims to extend the frameshift suppression concepts established in model enzymes to a clinically relevant phase-variable methyltransferase in *M. bovis* and to clarify the potential contribution of frameshift suppression to phenotypic heterogeneity.

## MATERIALS AND METHODS

### Plasmids and cloning of (AG)_n_ repeat variants

To systematically interrogate frameshift suppression in both the +1 and -1 translational frames, we engineered a series of Mod1’(0)::mEGFP translational fusion constructs, with Frame 0 as the in-frame no repeat control. These constructs were cloned into two expression backbones to enable differential transcriptional control: pHis17, which drives high-level expression under the T7 promoter, and pBR322, which enables expression under the native promoter.

For the pHis17 system, we generated a panel of (AG)_n_ repeat variants to assess repeat length-dependent modulation of translational frame restoration. The primers used for cloning these constructs are listed in the Supplementary File (Table S1). Frame +1 constructs were engineered with n= 2, 5, 8, 11, and 14 repeat units, whereas Frame -1 constructs comprised n= 3, 6, 9, 12, and 24 repeats. To establish a control for repeat-dependent effects at the PV SSR locus, the canonical (AG)_2_ repeat was disrupted by substituting it with the non-iterative sequence (AGAC). pSK (Empty) was used as an empty control, along with untransformed *E. coli* BL21(DE3) or *E. coli* BW25113, for baseline autofluorescence measurements.

For native promoter-driven expression in *E. coli* BW25113 using pBR322, a focused subset of repeat variants was selected to evaluate translational behaviour at physiological expression levels. Frame +1 constructs included n= 2, 11, and 14 repeats, while Frame -1 was represented by n= 9. The primers used for cloning these constructs are listed in the Supplementary File (Table S1). Sequences were confirmed with Sanger sequencing. This enabled a comparative analysis of high-expression (T7) and native-expression regimes while minimizing construct redundancy.

### Strains and growth conditions

We used *E. coli* BL21(DE3) strains (*lon-11, Δ(ompT-nfrA)885, Δ(galM-ybhJ)884, λDE3 [lacI, lacUV5-T7 gene 1, ind1, sam7, nin5], Δ46, [mal+]K-12(λS), hsdS10*) for the transformation and expression of the gene under T7 promoter (for pHis17 constructs). While for testing native expression (for pBR322 constructs), we used *E. coli* BW25113 *(F^-^ LAM^-^ rrnB3 DElacZ4787 hsdR514 DE(araBAD)567 DE(rhaBAD)568 rph-1).* Cultures were grown at 37°C in Miller Luria Bertani Broth (LB) supplemented with 100 µg/ml Ampicillin. Cultures were induced with IPTG (Isopropyl β-D-1-thiogalactopyranoside) at a 0.5-0.6 O.D._600_ and grown for 5 hours post-induction. Induction was performed on the secondary culture grown with pHis17, while cultures grown with pBR322 were incubated for 8 hours before harvesting the cells.

### Flow cytometry data acquisition

#### Sample Preparation

A 3ml aliquot of the induced culture was harvested by centrifugation, and the cell pellet was washed thrice with 1x Phosphate-Buffered Saline (PBS) (137 mM NaCl, 2.7 mM KCl, 10 mM Na_2_HPO_4, 1.8_ mM KH_2_PO_4_). Cells were fixed in 1 ml of 4% (w/v) paraformaldehyde for 15 min at 25°C, washed 3 times with 1x PBS, and diluted the sample to an O.D._600_ of 0.025 in 1x PBS. All steps mentioned above were performed on ice or at 4°C, unless otherwise specified.

#### Data acquisition

Flow cytometric acquisition was performed on a BD FACSCelesta^TM^ system using the ‘Medium’ flow rate setting (35 µL/min) with a forward scatter (FSC) threshold of 500. Detector voltages were set to 297 V for FSC, 294 V for side scatter (SSC), and 449 V for the FITC (Fluorescein Isothiocyanate) fluorescence channel. Events were acquired at approximately 100–200 events per second, and 10,000 events were collected per sample.

#### Gating strategy and statistical analysis

Flow cytometry data were analyzed using FlowJo™ software (version 10.9.0; BD Biosciences). An initial gate (P1) was defined on a bivariate density plot of side scatter area (SSC-A) versus forward scatter area (FSC-A) to delineate the primary cell population from a total of 10,000 acquired events. Samples in which the P1 population comprised less than 90% of total events were excluded from downstream analysis. The P1-gated population was further examined using log_10_-scaled univariate histograms of FITC fluorescence intensity. To distinguish fluorescence shifts attributable to differences in protein expression from those due to changes in cellular complexity, quadrant gating (Q1-Q4) was applied to SSC-A versus FITC-A density plots. Detailed statistical analysis of univariate FITC-A histograms was performed using OriginPro 2025 (Windows 11, version 10.2.0.196) after exporting the raw P1-gated population in .fcs format.

#### Calculation of noise (η) and mean of median fluorescence intensity 〈MFI⟩

Noise (η) in gene expression was quantified using the squared coefficient of variation (CV^2^), defined as the ratio of variance (σ^2^) to the square of population mean (µ^2^) as shown in the equation below. This definition follows the standard formulation used to quantify stochastic fluctuations in gene expression levels in single-cell studies (Elowitz et al., 2002; Swain et al., 2002). The mean 〈MFI⟩ was calculated by averaging the median fluorescence intensity from three independent biological replicates.

### Immunoblot sample preparation and analysis

After induction with 0.5 mM IPTG for 5 hours, 2.4 × 10^9^ cells were harvested and lysed in 100 µl of protein loading buffer (8% (w/v) SDS, 40% (v/v) glycerol, 0.25 M Tris-HCl pH 6.8, 10% (v/v) β-mercaptoethanol and bromophenol blue) by heating at 95°C for 15 minutes. Lysates were clarified by centrifugation at 20,000 rcf for 10 minutes at 4°C. Equal volumes (20 µl) of each sample were resolved on 12% SDS-PAGE at 120 V. Two identical gels were run in parallel: one processed for immunoblotting and the other stained with Coomassie Brilliant Blue to verify equal loading. Proteins were transferred to methanol-activated 0.22 µm PVDF membranes (HiMedia) using a wet transfer system (BioRad) at 90 V for 3 hours at 4°C in transfer buffer (25 mM Tris Base, 200 mM glycine, 20% methanol). Membranes were blocked for 1 hour at room temperature in 5% (w/v) non-fat skimmed milk prepared in TBST buffer (20 mM Tris-HCl pH 7.5, 150 mM NaCl, 0.1% (v/v) Tween-20). Blocked membranes were washed 3 times with 1x TBS buffer (20 mM Tris-HCl pH 7.5, 150 mM NaCl) for 5 minutes each. Blocked membranes were incubated at room temperature for 1 hour with primary antibody Anti-GFP Mouse mAb IgG2b (CST) diluted (1:20,000) in blocking buffer. After three washes with TBST buffer (5 minutes), membranes were incubated with HRP-conjugated Goat anti-Rabbit IgG (H+L) secondary antibody (Invitrogen) for 1 hour at room temperature. Following an additional 1x TBS wash, antibody binding was detected using enhanced chemiluminescence (Clarity western ECL, Bio-Rad). Signals were captured using the Amersham ImageQuant 800 (Cytiva) imaging system. The exposure time for the 1AG sample was 1 minute, while the rest of the samples had a 10-minute exposure time.

### Purification of GST-tagged Anti-GFP Nanobody (GST::GFP-Nb)

#### Plasmid, bacterial strain and growth conditions

The plasmid construct pGEX-6P1-GFP Nanobody was used for the purification of GST-tagged Anti-GFP nanobody (GST::GFP-Nb). For recombinant protein purification, 1 litre culture from (1:100) overnight starter culture with 100 µg/ml ampicillin as a selection marker was grown at 37°C at 200 rpm. Cells were induced with 0.2 mM IPTG at O.D._600_ **∼** 0.6. Post-induction culture was grown for 16 h at 18°C with shaking at 200 rpm. Cells were harvested, and the cell pellet was stored at -80°C.

#### Glutathione-based Affinity Batch purification protocol

Cell pellet (∼ 9 grams) was resuspended in 30 ml lysis buffer (10 mM Tris-HCl pH 7.5, 150 mM NaCl, 1mM EDTA, 1mM PMSF). Resuspended pellet was sonicated with 2 rounds of ultrasonication (60% amplitude, 3 min total sonication time, 1 s ON / 3 s OFF cycles). Crude lysate was clarified by centrifugation at 16,500 rpm for 30 min at 4°C. 5 ml slurry of Glutathione Sepharose 4B (GE Healthcare, Catalogue No: 17-0756-01 10 ml) was washed with 25 ml Binding buffer (10 mM Tris-HCl pH 7.5, 150 mM NaCl, 1 mM EDTA) at 800 rcf. 30 ml of cleared lysate was added to the beads and incubated for 4 hours at 4°C on a rolling shaker. Beads were washed 3 times with 25 ml Binding buffer by inverting, mixing, and spinning at 800 rcf for 5 min. Elution was carried out with Elution buffer (10 mM Tris-HCl pH 7.5, 150 mM NaCl, 1 mM EDTA, 10 mM reduced L-Glutathione). Five fractions of 1 ml were collected by gently shaking the beads with the elution buffer for 10 min. Fractions were resolved on 12% SDS-PAGE gel (41 kDa) and later concentrated with Amicon Ultra Centrifugal Filters 10K (Merck Millipore Ltd). Aliquots of 500 µl volume and a final concentration of 2.17 mg/ml were stored in -80°C. A gel image of the purified GST::GFP-Nb is shown in the Supplementary File (Figure S1). Before proceeding with the downstream purification of full-length protein from +1 frameshifted construct, a 500 µl aliquot of GST::GFP-Nb was diluted with 500 µl Binding buffer (total volume 1 ml) and dialyzed in 1 L Binding buffer overnight at 4°C to remove reduced L-glutathione.

### Purification of Full-length protein expressed from +1 frameshifted construct

#### Bacterial strain and growth conditions

To assess frameshift suppression and obtain the full-length protein from the +1 frameshifted construct bearing (AG)_2_ repeat insertion, it was expressed under the T7 promoter in *E. coli* BL21(DE3). An overnight starter culture grown at 37°C in LB media supplemented with ampicillin (100 µg/ml) was used to inoculate 4 L of secondary culture at a 1:100 ratio. Cultures were induced with 0.5 mM IPTG and harvested 5 h post-induction at 37°C. Cell pellets were resuspended in lysis buffer (10 mM Tris-HCl pH 7.5, 150 mM NaCl, 1 mM EDTA, 1 mM PMSF) and disrupted by 2 rounds of ultrasonication (60% amplitude, 3 min total sonication time, 1 s ON / 3 s OFF cycles). Lysates were clarified by centrifugation at 16,500 rpm for 30 min at 4°C.

#### GST::GFP-Nb based affinity capture

For affinity purification, 2 ml Glutathione Sepharose 4B resin (GE Healthcare, Catalogue No: 17-0756-01 10 ml) was equilibrated in the binding buffer (10 mM Tris-HCl pH 7.5, 150 mM NaCl, 1 mM EDTA) and incubated with dialyzed GST-tagged anti-GFP nanobody for 2 h to generate nanobody-coupled beads. Cleared lysate was applied to this resin, followed by washing with binding buffer. Bound proteins were eluted with binding buffer containing 10 mM reduced L-glutathione. Fractions containing the ∼40 kDa protein were dialyzed overnight at 4°C in binding buffer in the presence of Precision protease to remove the GST tag. The resulting sample was concentrated and resolved on 12% SDS-PAGE, revealing a ∼40 kDa band on Coomassie Brilliant Blue staining.

### Bottom-up proteomics with LC-MS/MS

#### In-gel enzymatic digestion

Purified protein was resolved on a 12% SDS-PAGE and visualized by Coomassie Brilliant Blue staining. Protein bands corresponding to ∼40 kDa were excised into 1 mm^3^ gel pieces with a sterile scalpel and transferred to fresh 1.5 ml tubes. The gel pieces were destained by multiple washes with 100 µl 50% acetonitrile (ACN)/ 50 mM ammonium bicarbonate (NH_4_HCO_3_) until clear, followed by dehydration with 100% ACN for 5 min to shrink the pieces. Proteins were reduced by incubating the gel pieces in 70 µl of 10 mM dithiothreitol (DTT)/ 50 mM NH_4_HCO_3_ for 1 h at 65°C. Samples were dehydrated again prior to alkylation. Alkylation was performed by incubating the gel pieces with 70 µl of 20 mM iodoacetamide (IAA) at 25°C for 20 min in the dark. Following alkylation, the gels were dehydrated as before. In-gel digestion was performed by adding 100 µL of digestion buffer containing sequencing-grade chymotrypsin (final concentration: 1 µg in 50 mM NH_4_HCO3; enzyme stock prepared at 0.5 µg/µL in 1 mM HCl). The samples were incubated overnight at 37°C to allow complete proteolysis.

#### Peptide extraction

Following digestion, peptides were sequentially extracted by adding 100 µL of extraction buffer containing 20% ACN/50 mM NH_4_HCO_3_, then 50% ACN/50 mM NH_4_HCO_3_, and finally 100% ACN. Each gradient of extraction buffer was incubated for 15 min with vigorous shaking at 25°C to maximize peptide recovery. The supernatant was collected and pooled after each gradient in a fresh 1.5 ml tube. Pooled extracts were vacuum-dried at 30°C in a SpeedVac. The resulting peptides were reconstituted in 80 µL of 0.1% trifluoroacetic acid (TFA) prior to desalting.

#### C18 desalting

Peptide desalting was performed using custom-made C18 microcolumns prepared from solid-phase extraction (SPE) disks (EmporeTM). Disks were punctured into 1 mm diameter plugs and inserted into the narrow end of 200 µl pipette tips to generate micro-C18 zip Tips. The columns were activated with 100% ACN followed by equilibration with 0.1% TFA. The peptide sample, reconstituted in 0.1% TFA, was slowly dispensed through a C18 resin for binding. This was followed by 160 µl of 0.1% TFA wash to remove salts and other hydrophilic impurities. Bound peptides were eluted directly into fresh tubes with 70 µl of 60% ACN/0.1% TFA. The eluates were vacuum-dried and reconstituted in 13 µl of 0.1% formic acid (FA) for LC-MS/MS analysis.

#### LC method

Desalted peptides were analysed using a high-resolution TripleTOF 6600 (AB Sciex instruments) mass spectrometer coupled to an Ekspert Nano LC 400 system. 9 µl desalted peptides in 0.1% FA were resolved on an analytical ChromXP C18 column (3 µm, 120 Å, 3C18-CL, 75 µm x 15 cm) maintained at 25°C. Chromatographic separation was achieved using a linear gradient of solvent B (0.1% FA in acetonitrile) against solvent A (0.1% FA in water) at a flow rate of 300 nl/min. Linear gradient of solvent B was as follows: 5% to 30% over 100 min, 30% to 95% over 3 min, 95% for 7 min. The column was then re-equilibrated with 5% solvent B and 95% solvent A prior to the next injection.

#### Time-Of-Flight MS/MS method and data acquisition

Tandem mass spectrometry data were obtained with TripleTOF 6600. The mass spectrometer was operated in positive ion mode with a nanospray voltage of 2.6 kV and an interface heater temperature of 120°C. Data were acquired in Information Dependent Acquisition (IDA) mode with the following optimized parameters: precursor ion charge state range from +2 to +5, and only ion intensities >50 counts per second were selected for fragmentation. The instrument was set to switch precursors after 13 MS/MS spectra, using advanced IDA settings with dynamic background subtraction and dynamic collision energy enabled. Precursor ions were dynamically excluded for 15 seconds following 2 occurrences with an exclusion mass tolerance of 50 ppm and a 6 Da isolation window to avoid redundant selection. The mass defect filter was enabled to ensure precursor consistency, and the system was configured to never permanently exclude former target ions, allowing re-acquisition if the signal reappeared above threshold. Smart IDA filters were applied to enhance spectral quality, with a fragment intensity multiplier of 2 and a maximum accumulation time of 2 seconds per MS/MS event. Spectra were acquired with Analyst TF (v1.7.1) in (.wiff) and (.wiffscan) file formats. The samples were analysed in the Mass Spectrometry Facility, Department of Biology, IISER Pune.

#### Peptide identification and analysis

The generated spectra were analysed using ProteinPilot^TM^ Software (v5.0, AB Sciex) with the Paragon^TM^ algorithm for database searching and protein identification. Searches were performed against the custom FASTA file with chymotrypsin specificity, Carbamidomethylation (Cys) as a fixed and methionine oxidation as a variable modification. Results were filtered to a global false discovery rate (FDR) < 1% at a 99% confidence threshold. The identified peptides from (.mzid) files were further analysed using Skyline software (Windows version v25.1.0.237).

## RESULTS

### Frameshift suppression dynamics revealed by (AG)_n_ repeat insertions in *mod1’::mEGFP*

To investigate the impact of phase-variable simple sequence repeats (PV SSRs) on translational rescue *in vivo,* variable lengths of (AG)_n_ repeat pHis17 (T7 promoter) constructs were expressed in *E. coli* BL21(DE3). These insertions were designed to disrupt the native reading frame and generate constructs with either a +1 or a -1 frame shift (Refer to Figure 1). Frameshift suppression was quantified at the single-cell level using flow cytometry-based fluorescence measurements.

**Figure 1:**
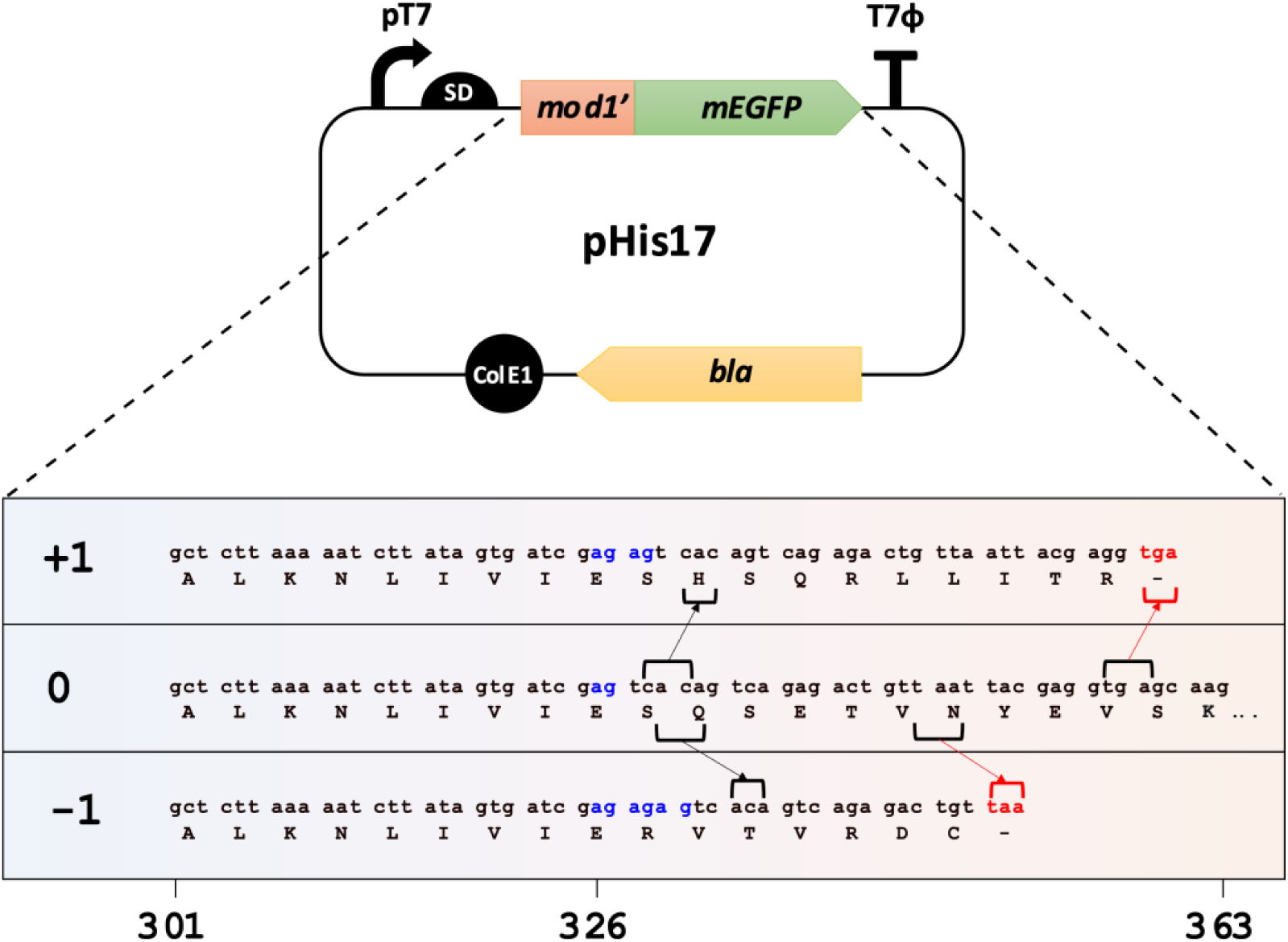
Nucleotide sequences translated into three distinct reading frames from Mod1’::mEGFP in pHis17 plasmid. Nucleotide sequences (5’→3’) shows forward translational frames of *mod1’* ORF (301-363): Frame 0 (IN-FRAME) and OUT-OF-FRAME (Frame +1 and -1). The schematic illustrates how insertion of (AG)_n_ repeat insertion (bold blue) enables frameshift and cause premature truncation at stop codons (bold red) unique to Frame +1 and -1.

#### Frame +1 constructs exhibit repeat-independent intermediate fluorescence

Flowcytometry analysis revealed that constructs with +1 frameshifts consistently produced an intermediate fluorescent population (light green populations) ∼10 fold higher than the basal autofluorescence level of the cells shown with ‘BL21(DE3)’ and ‘Empty’ control (grey populations) (Figure 2A and B). This indicated significant fluorescence from mEGFP expression in these constructs. Fluorescence intensity distributions across different (AG)_n_ repeat lengths (n = 2, 5, 8, 11, and 14) remained largely comparable. To check whether indels in the repeat region as small as 2AG were not causing frameshift suppression, we used a scrambled (AGAC) construct as the control. This scrambled control showed the same distribution profile as other +1 frameshifted samples. This suggested that the efficiency of +1 suppression was relatively independent of repeat length or even the presence of the repeat itself. The fluorescence signal observed in +1 Frame constructs was intermediate relative to the IN-FRAME (1AG) control, which indicated partial but reproducible rescue of PV SSR-mediated frameshifts. Populations followed a unimodal distribution across all samples. This unimodality across repeat variants suggested that +1 suppression originated from intrinsic errors introduced during protein expression. The possibilities for error included transcriptional slippage, ribosomal slippage, or an alternate translation initiation that would result in expression of mEGFP instead of Mod1’::mEGFP.

**Figure 2:**
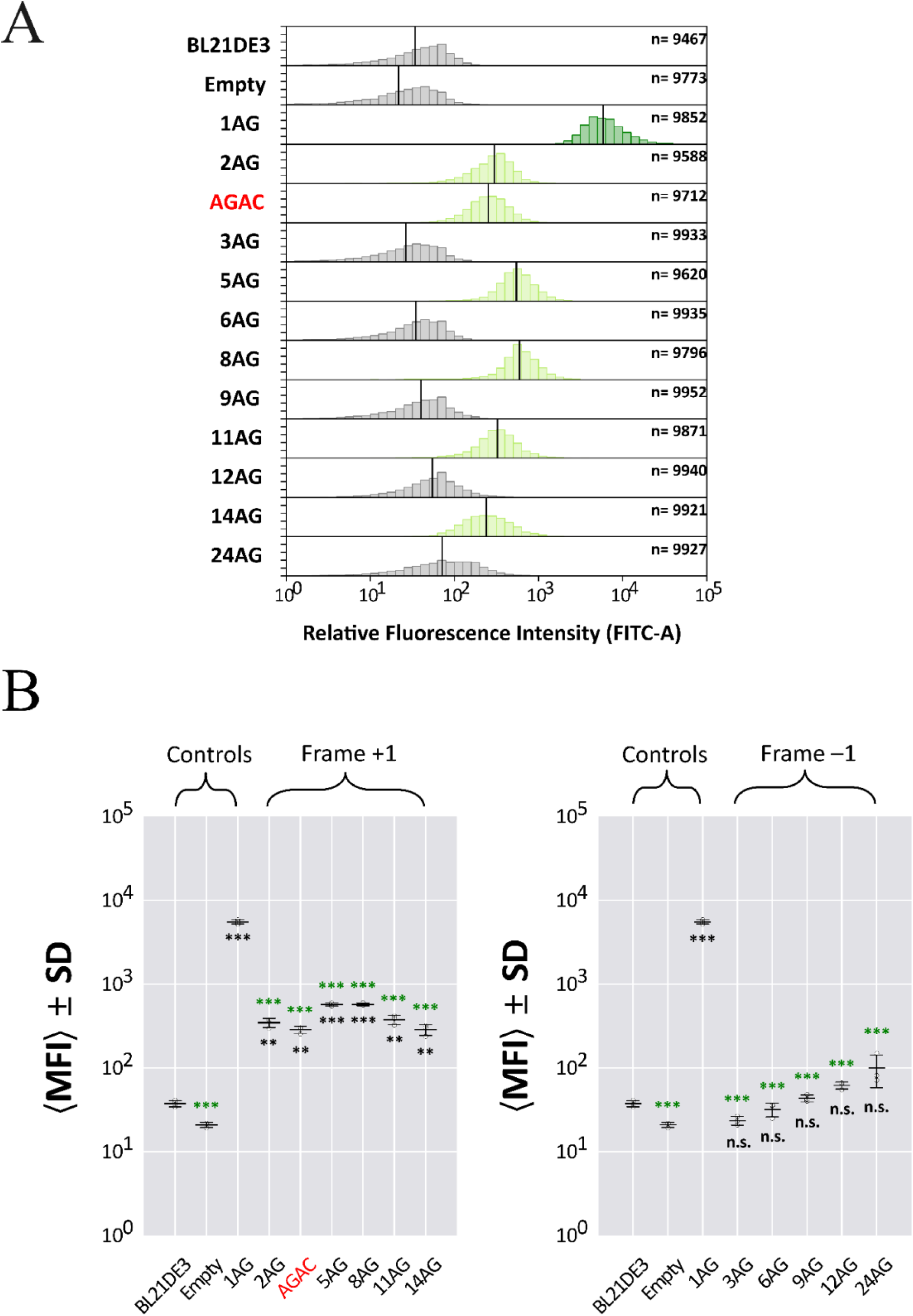
(AG)ₙ repeat tracts in *mod1’* modulate single-cell fluorescence levels. (A) Probability density histograms of *E. coli* BL21(DE3) cells expressing Mod1’::mEGFP with (AG)ₙ repeat insertions in the *mod1’* gene. Fluorescence was quantified via the FITC-A channel. +1 and -1 frameshifted constructs are shown in light green and grey histograms respectively. Positive control with in-frame (Frame 0) 1AG is shown in dark green. ‘BL21(DE3)’ and ‘Empty’ shown in grey are untransformed and empty negative controls respectively representing baseline autofluorescence. Vertical black lines mark the median fluorescence intensity (MFI). The number of events (cells) is indicated as ‘n’ in the top right corner of each histograms. The Y-axis depicts probability density (PD), calculated using the following formula: 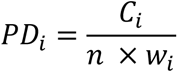 where *PD_i_* is the probability density for bin ‘i’, *C_i_* is the count of events in bin ‘i’, *n* is the total number of events, and *w_i_* = 0.1 is the bin width (constant). The Y-axis ranges from 0 to 2.5 with tick intervals at 0.5. The X-axis indicates log_10_-transformed relative fluorescence intensity. (B) Median fluorescence intensities (MFI) of Mod1’::mEGFP with +1 and -1 frameshifted (AG)_n_ repeats. Black horizontal line represents the mean of MFI from three independent biological replicates with error bars representing standard deviations (SD). Frame -1 (gray, right graph) and Frame +1 (light green, left graph) frameshifted constructs display distinct fluorescence profiles with increasing repeat lengths compared to in-frame 1AG as positive control and autofluorescence controls (BL21(DE3) and Empty as negative controls). Statistical comparisons were performed using one-way repeated-measures ANOVA. Fischer LSD post hoc comparison test was used to evaluate pairwise differences. A significant main effect was observed *(F(1, 2) = 6101.55255, p = 0.00016)*. Pairwise significance is denoted as follows: *p < 0.05 (*), p < 0.01 (**), p < 0.001 (***); n.s., not significant*. Pairwise significance symbols in black and dark green are in comparison with Empty and 1AG controls respectively.

#### Frame -1 suppression emerged gradually with increasing repeat length

In contrast, constructs with -1 frameshifts introduced with (AG)_n_ repeats (where n = 3, 6, 9, 12, and 24) displayed a different behaviour. Short (AG)_n_ repeats produced fluorescence levels comparable to the background autofluorescence, which indicated minimal frameshift suppression. However, as we increased the repeat length, a gradual emergence of a fluorescent population was evident, as the grey population shifted to the right in Figure 2A and B.

Despite this gradual shift in population towards higher fluorescence, fluorescence intensity in -1 constructs remained lower than that observed for +1 suppression across all repeat lengths tested. For -1 frameshifted constructs, suppression led to a more broadened population distribution and a weakly shifted peak, suggesting higher stochasticity in frameshift suppression events, as evidenced by the higher standard deviation observed with (AG)_24_ (Figure 2A and B).

#### Frameshift suppression established an expression vs. noise trade-off

In addition to mean expression output, we assessed population-level heterogeneity by calculating gene expression noise (η) as the squared coefficient of variation (CV^2^) of fluorescence intensity distributions. This metric enabled comparison of direct variability normalized to expression magnitude. This is widely used to characterize stochastic gene expression states in microbial populations (Elowitz et al., 2002). See Figure 3.

**Figure 3:**
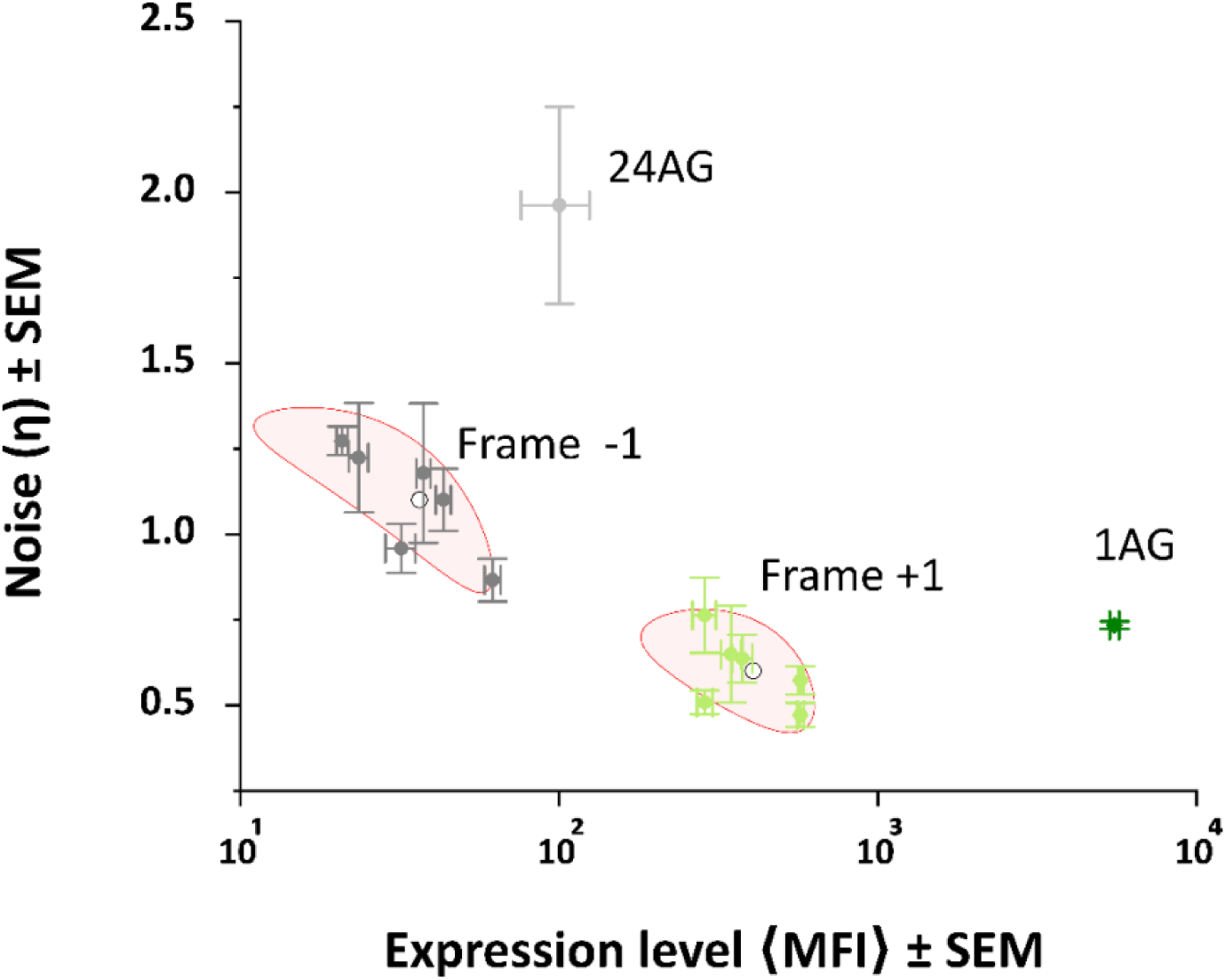
Translation in alternative reading frames imposes defined trade-offs between protein abundance and expression variability in a population. Noise in gene expression ‘η’ (measured as mean coefficient of variation squared 〈CV^2^⟩ on Y-axis) is plotted against mean of Median Fluorescence Intensity 〈MFI⟩ (Expression level on X-axis). Data points (solid circle) and error bars in the scatter plot represent the mean ± SEM (standard error of mean) for both the variables from the values obtained from three independent biological replicates (N=3). Clustering was performed with unsupervised k-means clustering (k=4) after performing *z-score* normalization. Clusters are shown as red ellipses with 95% confidence intervals and open circles depicts centroid of the cluster. The graph reveals that constructs with different reading frames segregate into distinct clusters (except 1AG and 24AG), showing that translational frameshifts not only shift expression levels but also reshapes the noise profile representing heterogeneity in the population.

Constructs with -1 frameshift suppression displayed markedly elevated noise relative to +1 constructs. Flow cytometry distributions showed broad fluorescence profiles with increased dispersion among individual cells, reflected in higher noise (η) values. Although longer (AG)_n_ repeats enabled detectable fluorescence, expression remained heterogeneous, with subpopulations exhibiting variable protein output. In the extreme case, (AG)_24_ showed the highest level of heterogeneity. In contrast, +1 frameshift suppression resulted in comparatively narrow fluorescence distributions and significantly lower noise (η) values. Despite producing only intermediate fluorescence relative to the in-frame control (AG)_1_, the +1 constructs maintained consistent expression across the population, indicating lower population heterogeneity. See Figure 3.

Overall, these results demonstrated a distinct population-level expression architecture driven by frameshift suppression. +1 suppression generated a *moderate-expression, low-noise state*, whereas -1 suppression produced a *low-expression, high-noise state* characterized by increased heterogeneity. Frameshift suppression in the presence of (AG)_n_ repeats acted as a regulatory layer that simultaneously modulated expression amplitude and population heterogeneity.

### An upstream (A)_5_T motif as an additional switch for the frameshift-rescue of +1 Frames

To validate whether fluorescence in +1 and -1 Frame (AG)_n_ constructs originated from expression of full-length Mod1’::mEGFP or alternative translation product (mEGFP only), protein outputs were analysed by immunoblotting and mass spectrometry.

Whole-cell lysates from frameshifted constructs that exhibited fluorescence were subjected to SDS-PAGE and immunoblotting with anti-GFP antibodies. For details, see Materials and Methods. Immunoblot analysis revealed a clear protein band corresponding to the expected molecular weight of full-length Mod1’::mEGFP (40 kDa) for all +1 Frame constructs. Mod1’::mEGFP expression got more prominent with longer repeats for -1 Frames, i.e., (AG)_9_, (AG)_12_, and (AG)_24_, with a visible shift for (AG)_24_ due to higher molecular weight compared to the rest among Frame -1. Refer to Figure 4A and B. In neither of the blots could we see a band corresponding to just mEGFP (∼27 kDa) indicating the absence of an alternate translation site. As a loading control for both the blots, we ran SDS-PAGE gels with the same amount of sample loads (refer to Supplementary File, Figure S2). Thus, fluorescence detected by flow cytometry can be directly related to full-length protein expression rather than translational bypass or reporter artifact.

**Figure 4:**
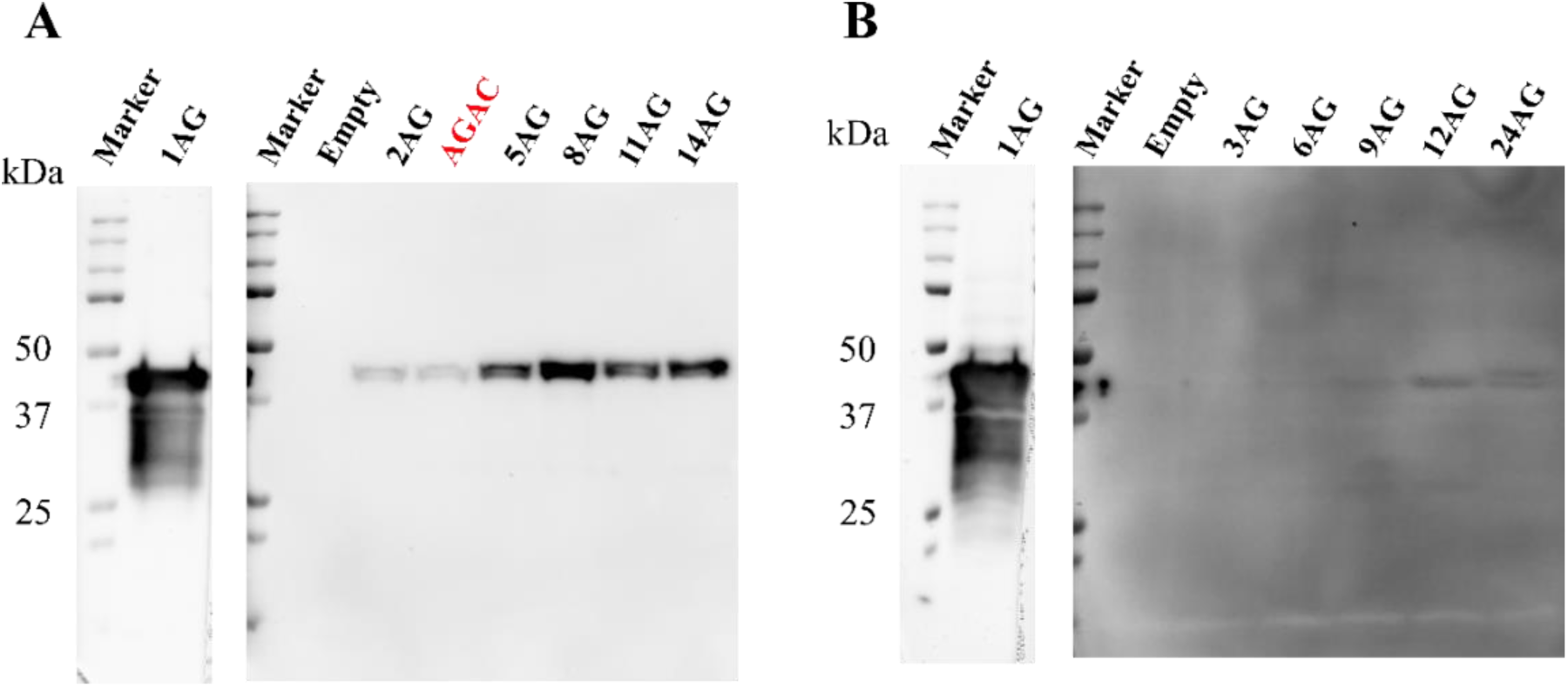
Translational frameshift suppression restores full-length protein expression. Immunoblot analysis of Mod1′::mEGFP (∼40 kDa) expression in +1 Frame (A) and -1 Frame (B) frameshifted constructs with variable lengths of (AG)ₙ repeats. “1AG” and “Empty” represent positive and negative controls, respectively. In both cases, the ∼40 kDa band indicates restoration of full-length protein due to frameshift suppression. There are no bands at ∼27 kDa for mEGFP. Blots were probed with the Mouse monoclonal anti-GFP antibody and the corresponding SDS-PAGE gels stained with Coomassie Brilliant Blue is shown in ‘Supplementary File, Figure S2’ to show equal loading across samples (See **‘**Materials and Methods’ section).

#### Proteomic identification of the frameshift-derived peptide confirmed -1 compensatory frameshift at (A)_5_T motif rescued +1 Frames

To pinpoint the region associated with frame recovery, affinity-purified Mod1’::mEGFP was subjected to chymotrypsin digestion followed by mass spectrometric analysis. For details, refer to Materials and Methods and Figure 5A, B, and C. For this analysis, we purified the full-length protein from the +1 frameshifted (AG)_2_ construct. Peptide mapping via a bottom-up mass spectrometry approach confirmed extensive coverage of the wild-type Mod1’::mEGFP sequence, as shown by the green-highlighted sequence in Figure 5D. Peptides spanning the interval between the (A)5T tract and the (AG)2 repeats exhibited altered sequence signatures, revealing a short, *frameshift-derived peptide segment (SYSDR).* Downstream peptide fragments reverted to the expected wild-type sequence, indicating restoration of the original reading frame *(ESQSET…)* prior to completion of translation. Precursor peptide ion *SDRESQSETVNY* (Charge 2+) with observed *m/z* 707.8077 was identified at 32.85 min retention time as shown in the Extracted Ion Chromatogram (XIC) in Figure 5D. Fragmentation pattern of this precursor peptide ion was confirmed with identification of *b-* and *y-*ions in +1 and +2 charge states in MS/MS spectra (Figure 5E-F; Supplementary Table S2).

**Figure 5:**
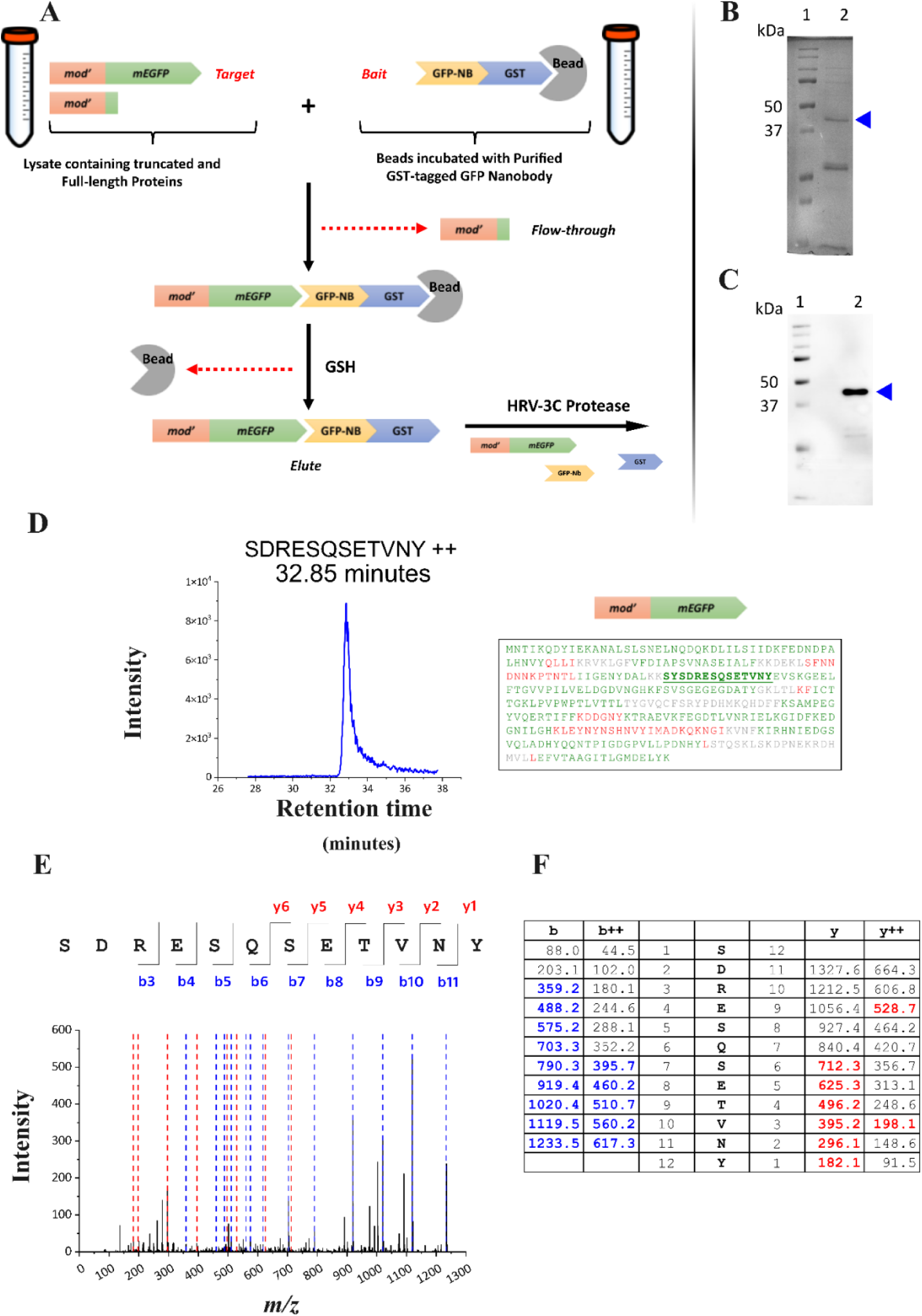
Purification and mass spectrometric validation of the frameshift-suppressed protein: (A) Schematic representation of GFP-Nb based affinity capture of full-length Mod1’::mEGFP translated from Frame +1 frameshifted (AG)_2_ construct (see ‘Materials and methods’ for details). (B) Coomassie stained SDS-PAGE analysis post purification using GFP-Nb conjugated beads. A prominent band corresponding to the full-length protein (∼40 kDa; blue triangle, lane 2) was excised and subjected to in-gel digestion followed by LC-MS/MS peptide identification. Lane 1 shows the molecular weight marker. (C) Immunoblot analysis of the purified sample probed with Mouse monoclonal anti-GFP primary antibody, confirming the identity of the ∼40 kDa full-length Mod1’::mEGFP protein (blue triangle, Lane 2). Lane 1 shows the molecular weight marker. (D) Extracted ion chromatogram (XIC, left) showing the *frameshift-derived peptide* detected in the chymotryptic digest of the purified protein. The peptide *SDRESQSETVNY* (charge 2+) was observed at *m/z* 707.8077 (charge 2+) with a retention time of 32.85 min. The right panel depicts the ProteinPilot-generated sequence coverage map of the full-length protein, with peptides colored by spectral confidence: high (green), moderate (yellow), low (red), and regions lacking spectral evidence (gray). The *frameshift-derived peptide (SYSDR)* followed by wild-type peptide is highlighted in bold green and underlined. (E) MS/MS fragmentation spectrum validating the identity of the frameshift-derived peptide. Collision-induced dissociation of the precursor ion shown in (D) produced *b-* and *y-*ions consistent with the theoretical fragmentation pattern. The left panel shows the MS2 spectrum annotated with *b-*ions (blue) and *y-*ions (red). The table (right) lists the observed *m/z* values for *b-* and *y-*ions in +1 and +2 charge states, with matched ions highlighted in blue and red and unmatched ions in gray.

These results demonstrated that +1 frameshifted constructs were rescued by the errors introduced in the upstream (A)_5_T tract, as shown in the black box in Figure 6. We speculate either of the two possibilities which might lead to Frame +1 rescue: (A) Transcriptional slippage by T7 RNA polymerase in *E. coli* BL21(DE3) leading to (+1A) insertion in the (A)_5_T tract, which could lead to -1 compensatory frameshift, thus causing frameshift suppression for +1 Frame. (B) Ribosomal Frameshift at (A)_5_T motif causing -1 frameshift rescuing +1 Frame constructs.

**Figure 6:**
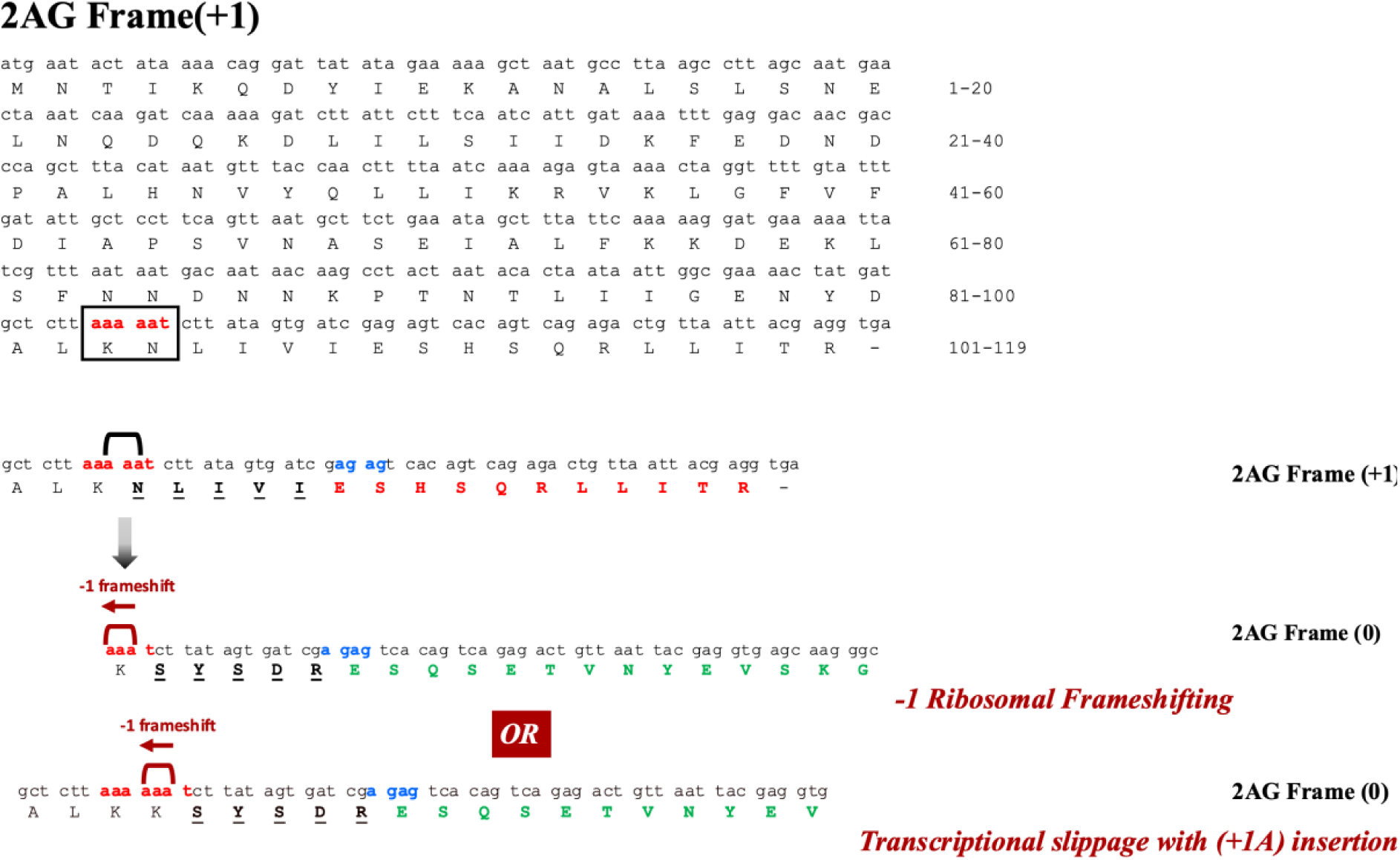
Translational reading frame of (AG)_2_ +1 Frame construct. Sequences show (AG)_2_ Frame (+1) construct translates (*NLIVI)* (black underlined) at the region between (A)_5_T and (AG)_2_ followed by nonsense peptide (bold red) terminating at TGA stop codon. (A)_5_T can rescue this frame via compensatory -1 frameshift either by: (i) -1 Ribosomal Frameshift, or (ii) Insertion of +1A due to transcriptional slippage. These two events bring back this frame to Frame (0) with the signature *frameshift-derived peptide (SYSDR)* between (A)_5_T and (AG)_2_.

### Native promoter expression attenuates the rescue of +1 Frame

To determine whether frameshift suppression efficiency depended on gene expression level, Frame +1 and Frame -1 (AG)_n_ constructs were expressed from the pBR322 vector under its native promoter in the *E. coli* BW25113 strain, rather than the high-expression T7-driven system used previously. This comparison enabled evaluation of how reduced transcriptional and/or translational flux influenced the restoration of protein expression. We chose limited (AG)_n_ constructs to test this: n=2, 11, and 14 for Frame +1, and n=9 for Frame -1.

Under native promoter expression, Frame +1 constructs exhibited a marked reduction in fluorescence intensity relative to their counterparts expressed from the T7 promoter. Flow cytometry histograms showed a substantial shift of population distributions towards lower fluorescence, indicating decreased production of functional Mod1’::mEGFP.

The intermediate fluorescence phenotype characteristic of Frame +1 suppression under strong expression conditions (T7 promoter) was no longer observed under native promoter expression. Instead, under native promoter expression fluorescence levels approached those previously measured for Frame-1 constructs (Figure 7A and B). This reduction indicated that efficient Frame +1 suppression was sensitive to expression strength and was significantly diminished when transcriptional output was lowered. Immunoblot analysis showed no band for Mod1’::mEGFP for Frame +1 and -1. See Figure 7C.

**Figure 7:**
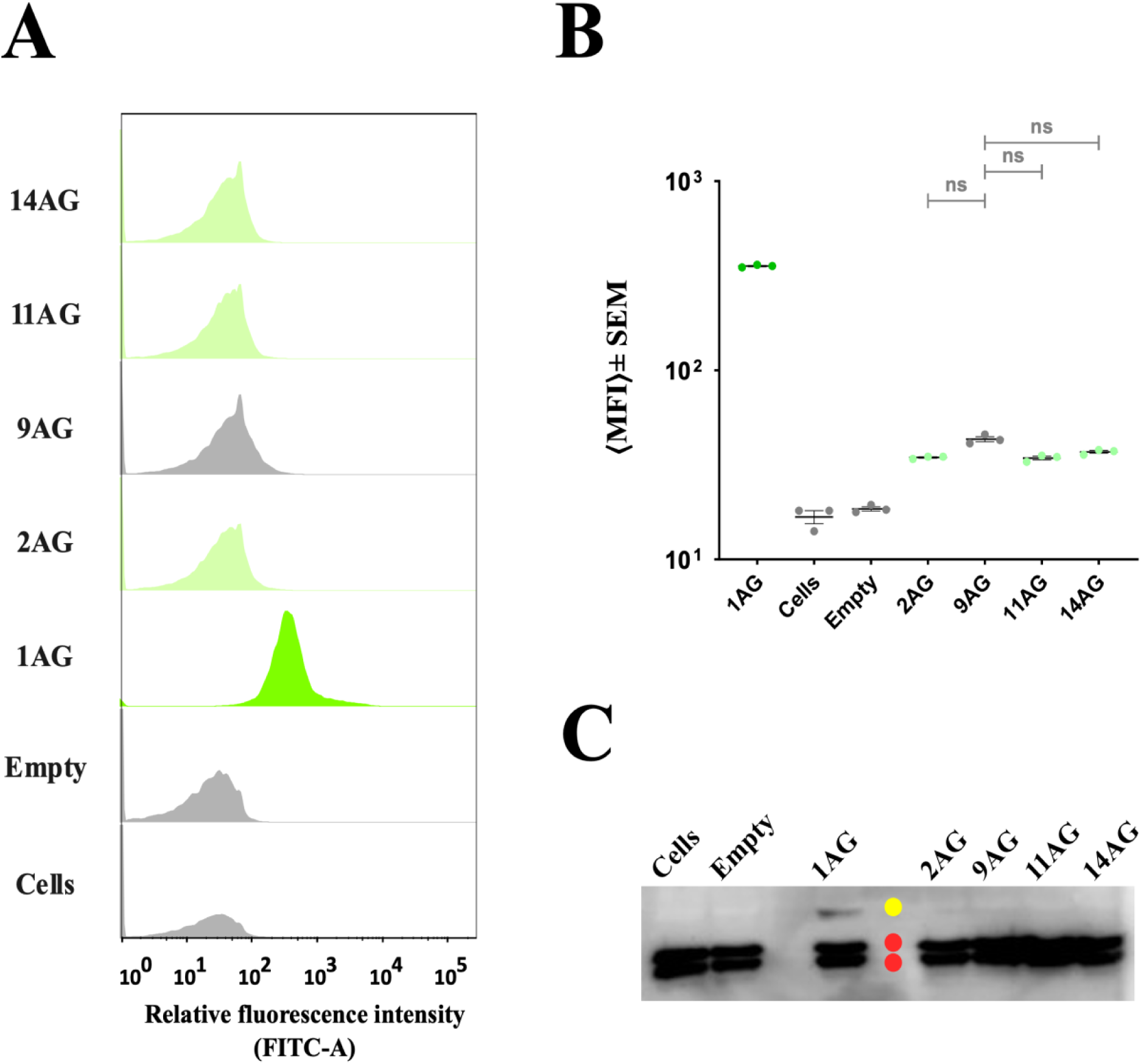
Frame +1 frameshift suppression reduces under lower transcriptional load. (A) Flowcytometry histograms of *E. coli* BW25113 cells expressing Mod1’::mEGFP under native promoter in pBR322 plasmid with (AG)ₙ repeat insertions in the *mod1’* gene. Fluorescence was quantified via the FITC-A channel. -1 and +1 frameshifted constructs are shown in light green and grey histograms respectively. Positive control in-frame (Frame 0) 1AG is shown in dark green. ‘Cells’ and ‘Empty’ shown in grey are untransformed and empty negative controls respectively representing baseline autofluorescence. (B) Graph showing mean of Median Fluorescence Intensity ⟨MFI⟩ from 3 independent biological replicates (N=3). Error bars represent standard error of mean (SEM). Statistical comparisons were performed using one-way repeated-measures Brown-Forsythe ANOVA test. Tamhane’s T2 post hoc comparison test was used to evaluate pairwise differences. A significant main effect was observed *(F* (DFn, DFd = 7071 (6.000, 4.133), p = <0.0001).* Pairwise significance is denoted as follows: *p < 0.05 (*), p < 0.01 (**), p < 0.001 (***); n.s., not significant*. Pairwise significance symbols in gray ‘ns’are in comparison with Frame -1 (9AG). (C) Immunoblot probed with polyclonal rabbit anti-GFP primary antibody. Yellow circle shows the band corresponding to full-length Mod1’::mEGFP in (1AG) and red circles are the non-specific bands of endogenous proteins of *E. coli* BW25113.

In conclusion, these findings established promoter context as a key determinant for frameshift suppression outcomes. While sequence elements within the ORF enabled restoration of protein expression, the extent to which this restoration manifested at the population level depended on the transcriptional strength.

## DISCUSSION

### Phase-variable SSRs in Type III *mod* ORF tunes translational outcomes

Phase-variable simple sequence repeats (PV SSRs) are traditionally studied as drivers of genetic phase variation at the DNA level for type III phasevarions (Nahar et al., 2023; Srikhanta et al., 2009, 2005). However, the present study demonstrates that repeat elements embedded within coding sequences can also reshape protein expression outcomes at the translational level. By strategically introducing (AG)_n_ repeats into the *mod1’::mEGFP* open reading frame, we show that frameshift suppression generates distinct expression regimes characterized not only by differences in mean protein output but also by fundamentally different population behaviours.

A central finding of this study is the strong asymmetry between +1 and −1 frameshifted constructs. Frame +1 variants consistently produced *intermediate fluorescence independent of repeat length*, whereas -1 constructs required progressive repeat expansion before detectable expression emerged. Even under permissive conditions, -1 suppression never reached the expression levels observed for +1 constructs.

This *directional bias* suggests that the presence of the repeat does not solely determine the restoration of the translation frame. Previous studies have shown that frameshift outcomes are highly sensitive to local nucleotide composition and kinetic constraints during transcription and translation (Atkins et al., 1972; Farabaugh, 1996; Liao et al., 2011). The repeat-independent behaviour of +1 constructs observed here implies that correction events arise from intrinsic properties of the coding environment rather than repeat-length dependent structural effects alone. In contrast, the gradual increase in fluorescence upon increase of repeat number observed for −1 constructs indicates that longer repeats enhance the probability of frame restoration but do so inefficiently, producing heterogeneous expression states rather than uniform recovery.

Such genetic systems act as efficient information processing platforms that execute logical operations via molecular interactions (Bashor and Collins, 2018; Nielsen et al., 2016). In contrast to conventional synthetic circuits that rely on transcriptional regulators or recombinases, this study demonstrates that coding DNA sequences can directly encode logic gates. Our results indicate that the interaction between (AG)_n_ repeat-induced frame disruption and an upstream (A)₅T homo-polymeric tract generates a functional outcome equivalent to an *XNOR* logical operation at the level of protein synthesis. The truth table explaining the XNOR logic for expression is shown in Figure 8. Rather than acting independently, these two sequence elements collectively determine whether full-length Mod1’::mEGFP is produced and to what level.

**Figure 8:**
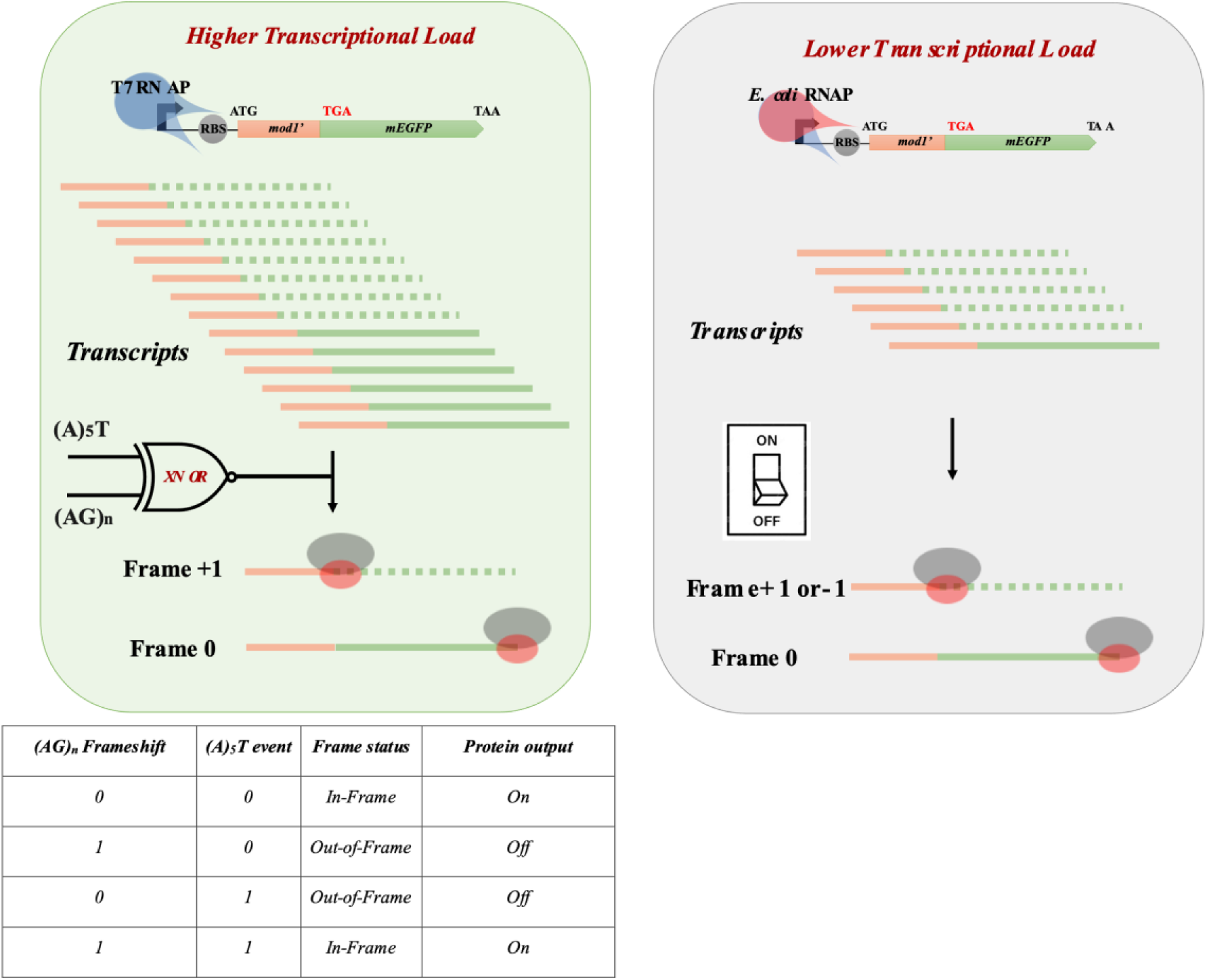
Model showing the Type III *mod* CDS following XNOR logic at higher transcriptional strength. The truth table explaining the XNOR logic gated expression is shown below. The same circuit turns into a binary ON/OFF switch at lower transcriptional strength.

### Promoter strength tuned logic gate fidelity for +1 Frames

Transition from T7-driven expression to native promoter control attenuated the frameshift suppression of Frame +1. This observation suggests that transcriptional flux can modulate the effective protein output of the Type III *mod* gene of *M. bovis.* This effect may be due to lower mRNA levels, which reduces the detection of frameshift suppressed products. It could also be caused by differences in transcriptional slippage between the *E. coli* RNA polymerase and the T7 RNA polymerase, or by ribosomal slippage during translation. Our study does not distinguish between these possibilities to identify the cause of the difference in the level of expression in frameshifted +1 Frame.

Higher transcriptional load increases the likelihood of transcriptional errors being introduced at (A)_5_T logical input for Frame +1. Reduced expression lowers this probability, collapsing the XNOR logic and converting the system into a *binary ON/OFF* switch. Expression-rate dependence has been observed across multiple stochastic gene expression systems, in which increased transcription amplifies rare molecular events into measurable phenotypes (Pedraza and Paulsson, 2008). Expression-dependent logic fidelity is present in engineered genetic circuits, in which signal amplitude determines switching reliability (Daniel et al., 2013). The present findings extend this principle to coding sequence-encoded logic. The generalized current model, as understood to date, is shown in Figure 8.

### Frameshift suppression leads to a distinct Expression-Noise landscape

Another important part of this study is the quantitative analysis of expression variability using the squared coefficient of variation (CV²) as a measure of gene expression noise (η). While many studies focus primarily on average protein levels, increasing evidence suggests that phenotypic variability itself represents a selectable biological trait (Elowitz et al., 2002; Raj and Van Oudenaarden, 2008). Here, +1 suppression generated a moderate-expression, low-noise regime, whereas -1 suppression established a low-expression, high-noise regime. This inverse relationship between expression magnitude and variability reflects a classical expression-noise trade-off, where inefficient production amplifies stochastic fluctuations (Swain et al., 2002; Taniguchi et al., 2010).

The unimodal distributions observed across +1 constructs further indicate population-wide engagement of the rescue mechanism. On the other hand, broadened distributions in -1 constructs reveal heterogeneous engagement across individual cells, suggesting that only a fraction of the population achieves productive expression at any given time.

From a synthetic biology perspective, these findings highlight frameshift suppression as a mechanism that can encode programmable heterogeneity without altering promoter architecture or requiring extra regulatory proteins. Such intrinsic noise modulation resembles engineered strategies used to diversify phenotypes in synthetic circuits designed for bet-hedging or division of labour (Ackermann, 2015).

## SUPPLEMENTARY DATA

Supplementary data is available in ‘Supporting File’ (PDF).

## AUTHOR INFORMATION

### Corresponding Author

**Kayarat Saikrishnan -** Department of Biology, Indian Institute of Science Education and Research, Pune 411008, India; https://orcid.org/0000-0003-4177-8508; Email ID

### Authors

**Rushik Bhatti -** Department of Biology, Indian Institute of Science Education and Research, Pune 411008, India; https://orcid.org/0000-0002-8601-394X

### Author Contributions

R.B. carried out cloning, flow cytometry, protein purification, western blot and mass spectrometry. R.B. carried out data analysis of flow cytometry and proteomics mass spectrometry. K.S. conceptualized the study and participated in data analysis. R.B. and K.S. wrote the manuscript.

### Notes

The authors declare no competing financial interest.

## ACKNOWLEDGEMENTS

R.B. and K.S. would like to thank Indian Council of Medical Research (ICMR) for Senior Research Fellowship. R.B. would like to thank Om Mahakal, Baishali Das, Somadatta Naskar, and Rajdip Sarkar for their contributions in cloning some of the constructs. R.B. and K.S. would like to thank Saurabh Vaishnav and the Flow Cytometry facility, IISER Pune, which is part of the Pune Biotech Cluster, and Shekh Saddam Husen and the Mass spectrometry facility, IISER Pune.

