## Supplementary data for "Emergence of a complex logic gate from the integration of slippage-induced frameshift mechanisms"

### Table S1: List of primer sequences used for cloning.

| Sr. No. | Primer name | | Primer sequence (5'-3') |
|---|---|---|---|
| 1 | R3_pHis17_mod1(0)350_EGFP | | GAACAGCTCCTCGCCCTTGCTCACCTCGTAATTAACAGTCTCTGACT |
| 2 | EGFP_R | | CTTGTACAGCTCGTCCATGCCGAGAG |
| 3 | F3_pHis17_mod1(0)350_EGFP | | ATTTTGTTTAACTTTAAGAAGGAGATATACATATGAATACTATAAAACAGGATTATATAG |
| 4 | F16_pHis17_mod1(2)352 | $(AG)_2$ | CTTAAAAATCTTATAGTGATCGAGAGTCACAGTCAGAGACTG |
| 5 | F15_pHis17_mod1(3)354 | $(AG)_3$ | CTTAAAAATCTTATAGTGATCGAGAGAGTCACAGTCAGAGACTG |
| 6 | F8_pHis17_mod1(5)358 | $(AG)_5$ | CTTAAAAATCTTATAGTGATCGAGAGAGAGAGTCACAGTCAGAGACTG |
| 7 | F13_pHis17_mod1(6)360 | $(AG)_6$ | CTTAAAAATCTTATAGTGATCGAGAGAGAGAGAGTCACAGTCAGAGACTG |
| 8 | F14_pHis17_mod1(8)364 | $(AG)_8$ | CTTAAAAATCTTATAGTGATCGAGAGAGAGAGAGAGAGTCACAGTCAGAGACTG |
| 9 | F9_pHis17_mod1(9)366 | $(AG)_9$ | CTTAAAAATCTTATAGTGATCGAGAGAGAGAGAGAGAGAGTCACAGTCAGAGACTG |
| 10 | F10_pHis17_mod1(11)370 | $(AG)_{11}$ | CTTAAAAATCTTATAGTGATCGAGAGAGAGAGAGAGAGAGAGAGTCACAGTCAGAGACTG |
| 11 | F4_pHis17_mod1(12)372 | $(AG)_{12}$ | CTTAAAAATCTTATAGTGATCGAGAGAGAGAGAGAGAGAGAGAGAGTCACAGTCAGAGACTG |
| 12 | F21_pHis17_mod(14) | $(AG)_{14}$ | AATCTTATAGTGATCGAGAGAGAGAGAGAGAGAGAGAGAGAGAGAGTCACAGTCAGAGACTG |
| 13 | pBR322_mod_F | Cloning | CTAACGCAGTCAGGCACCGTGTCATATGAATACTATAAAACAGGATTATATAGAAAAAG |
| 14 | pBR322_mod_R | Cloning | CACGATGCGTCCGGCGTAGAGGATCCTTACTTGTACAGCTC |

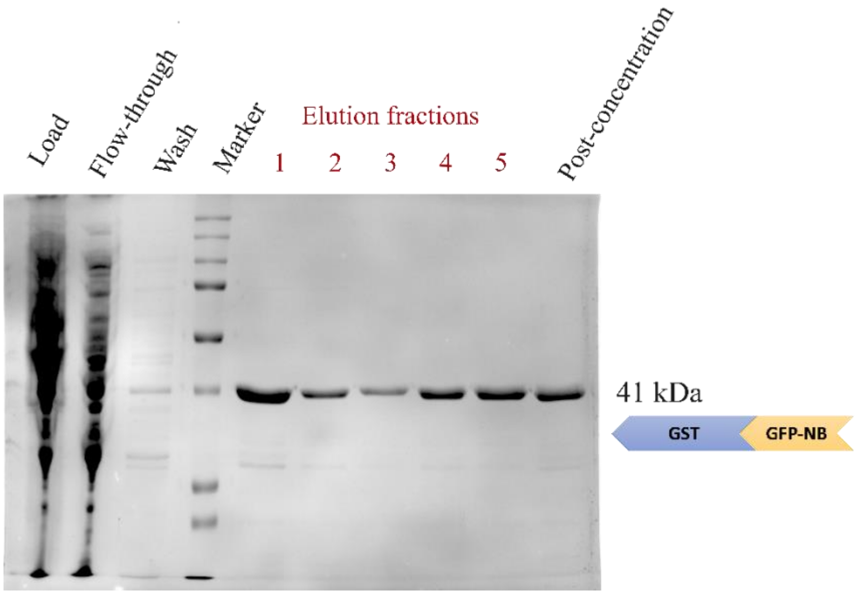


**Figure S1:** GST-tagged Anti-GFP Nanobody (GST::GFP-Nb) purification by Glutathione-based affinity capture. Molecular weight of GST::GFP-Nb is 41 kDa. Marker used is Precision Plus Protein$^{TM}$ Standards (10-250 kDa).

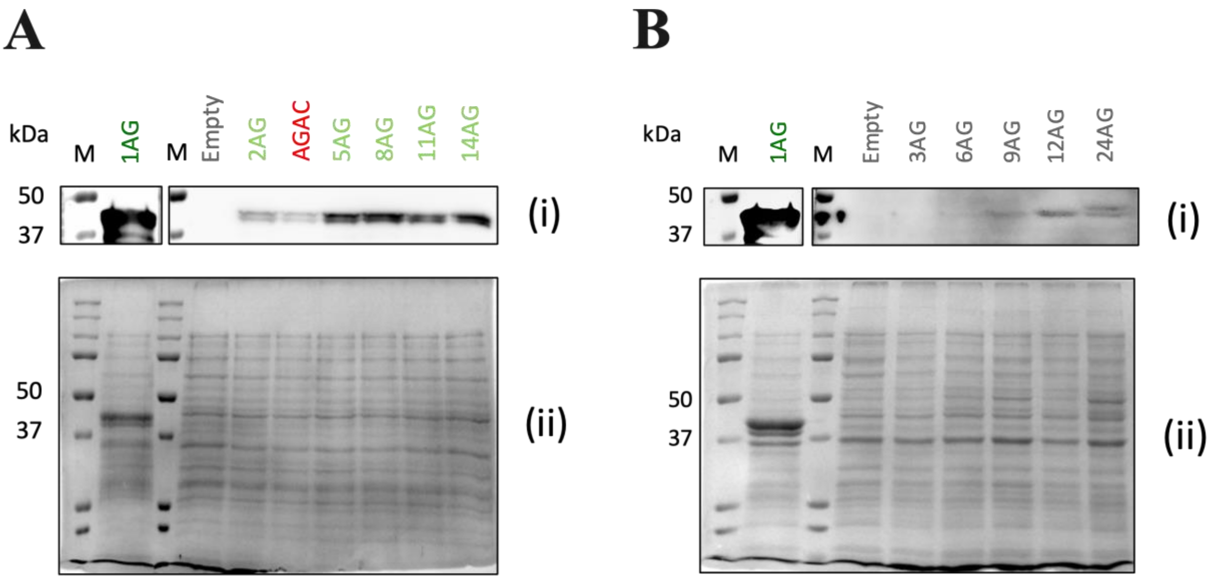


**Figure S2:** Translational frameshift suppression restores full-length protein expression. Western blot analysis of Mod1′::mEGFP (~40 kDa) expression in +1 Frame (A) and -1 Frame (B) frameshifted constructs containing variable lengths of $(AG)_n$ repeats. “1AG” and “Empty” represent positive and negative controls, respectively. In both cases, the reappearance of the ~40 kDa band indicates restoration of full-length protein upon frameshift suppression. Blots were probed with the monoclonal anti-GFP antibody (panels i), and corresponding SDS-PAGE gels stained with Coomassie Brilliant Blue (panels ii) confirm equal loading across samples (See ‘Materials and Methods’ section).

**Table S2: Distinct peptide summary from the ProteinPilot™ Software (v5.0, AB Sciex).**

This table summarizes peptide-level identifications and associated quantitative and acquisition parameters obtained from LC-MS/MS analysis. *Peptide Locus* indicates the protein origin of the identified peptide. *Best Conf (Peptide)* represents the highest confidence score assigned to the peptide spectrum match. *Best Hypoth Conf* denotes the confidence of the corresponding protein hypothesis. *Sequence* provides the amino acid sequence of the identified peptide. *Obs m/z* and *Theor m/z* correspond to the observed and theoretical mass-to-charge ratios, respectively. *Theor z* indicates the assigned charge state. *Acq Time* denotes the retention time at which the peptide was acquired. Quantitative measures include *Intensity (Peptide)*, which represents the integrated signal intensity across the peptide peak, and *Intensity at Acq*, which indicates the signal intensity at the acquisition time point. Chromatographic features include *Apex Time (Peptide)*, the time at maximum signal intensity, and *Elution Peak Width (Peptide)*, reflecting the duration of peptide elution. *Sum MS2 Counts* represents the total number of MS/MS spectra acquired for a peptide and serves as an indicator of sampling depth and identification robustness. Precursor peptide ion *SDRESQSETVNY* (Charge +2) is described in the Figure 5.

| Peptide Locus | Best Conf (Peptide) | Best Hypoth Conf | Sequence | Obs m/z | Theor m/z | Theor z | Acq Time | Intensity (Peptide) | Intensity at Acq | Apex Time (Peptide) | Elution Peak Width (Peptide) | Sum MS2 Counts |
|---|---|---|---|---|---|---|---|---|---|---|---|---|
| 1.001 | 99 | 99 | MNTIKQDYIEKANAL | 589.9696 | 589.9697 | 3 | 54.08272 | 18264.85 | 18072.83 | 54.06 | 0.3 | 1526.854 |
| 2.001 | 99 | 99 | MNTIKQDYIEKANAL | 595.2981 | 595.3013 | 3 | 54.04957 | 4996.61 | 4376.74 | 54.13 | 0.33 | 576.0717 |
| 2.002 | 99 | 98.56 | MNTIKQDYIEKANAL | 595.2981 | 595.3013 | 3 | 54.04957 | 4996.61 | 4376.74 | 54.13 | 0.33 | 576.0717 |
| 3.001 | 99 | 99 | MNTIKQDYIEKANAL | 590.2963 | 590.2977 | 3 | 53.4706 | 2947.24 | 2933.76 | 53.48 | 0.27 | 492.1873 |
| 3.002 | 99 | 98.56 | MNTIKQDYIEKANAL | 590.2959 | 590.2977 | 3 | 56.60842 | 3399.68 | 1962.83 | 56.77 | 0.44 | 345.8504 |
| 3.003 | 99 | 95.17 | MNTIKQDYIEKANAL | 590.2959 | 590.2977 | 3 | 56.57825 | 3399.68 | 1857.02 | 56.77 | 0.44 | 345.4136 |
| 4.001 | 99 | 99 | MNTIKQDYIEKANAL | 598.9722 | 598.9733 | 3 | 54.988 | 2706.37 | 2586.57 | 55.03 | 0.37 | 514.3929 |
| 4.002 | 99 | 99 | MNTIKQDYIEKANAL | 598.9722 | 598.9733 | 3 | 54.988 | 2706.37 | 2586.57 | 55.03 | 0.37 | 514.3929 |
| 4.003 | 99 | 95.17 | MNTIKQDYIEKANAL | 598.9722 | 598.9733 | 3 | 54.988 | 2706.37 | 2586.57 | 55.03 | 0.37 | 514.3929 |
| 5.001 | 22.9 | 22.91 | MNTIKQDYIEKANAL | 603.9721 | 603.9732 | 3 | 46.37117 | 2094.04 | 1836.7 | 46.43 | 0.2 | 1334.605 |
| 7.001 | 99 | 99 | NTIKQDYIEKANAL | 540.9542 | 540.9579 | 3 | 53.15911 | 4949.85 | 4906.19 | 53.15 | 0.24 | 386.0238 |
| 8.001 | 34.5 | 34.52 | KQDYIEKANAL | 646.8133 | 646.8459 | 2 | 106.4042 | 4389.87 | 1388.91 | 106.69 | 0.33 | 151.5647 |
| 9.001 | 99 | 99 | SLSNELNQDQKDL | 752.366 | 752.3679 | 2 | 56.28277 | 45840.27 | 45083.86 | 56.27 | 0.33 | 2956.672 |
| 10.001 | 99 | 99 | SLSNELNQDQKDL | 752.8558 | 752.8599 | 2 | 61.29182 | 8701.72 | 6834.28 | 61.37 | 0.36 | 500.8528 |
| 10.002 | 99 | 95.17 | SLSNELNQDQKDL | 752.8558 | 752.8599 | 2 | 61.29182 | 8701.72 | 6834.28 | 61.37 | 0.36 | 500.8528 |
| 11.001 | 99 | 99 | SLSNELNQDQKDL | 780.875 | 780.8787 | 2 | 56.15043 | 5622.34 | 5328.33 | 56.2 | 0.4 | 695.7019 |
| 11.002 | 99 | 99 | SLSNELNQDQKDL | 780.875 | 780.8787 | 2 | 56.15043 | 5622.34 | 5328.33 | 56.2 | 0.4 | 695.7019 |
| 11.003 | 99 | 95.17 | SLSNELNQDQKDL | 780.875 | 780.8787 | 2 | 56.15043 | 5622.34 | 5328.33 | 56.2 | 0.4 | 695.7019 |
| 12.001 | 99 | 99 | SLSNELNQDQKDL | 780.8764 | 780.8787 | 2 | 58.43597 | 4380.42 | 4295.77 | 58.42 | 0.35 | 311.6418 |
| 13.001 | 99 | 99 | SLSNELNQDQKDL | 752.8551 | 752.8599 | 2 | 57.10405 | 3769.08 | 3766.31 | 57.11 | 0.31 | 592.4759 |
| 13.002 | 99 | 99 | SLSNELNQDQKDL | 752.8551 | 752.8599 | 2 | 57.10405 | 3769.08 | 3766.31 | 57.11 | 0.31 | 592.4759 |
| 14.001 | 98.6 | 98.56 | SLSNELNQDQKDL | 760.3608 | 760.3597 | 2 | 56.1471 | 7055.72 | 3350.13 | 56.27 | 0.33 | 654.0914 |
| 14.002 | 98.6 | 98.56 | SLSNELNQDQKDL | 760.3608 | 760.3597 | 2 | 56.1471 | 7055.72 | 3350.13 | 56.27 | 0.33 | 654.0914 |
| 17.001 | 99 | 99 | NQDQKDLI | 487.2399 | 487.2511 | 2 | 68.15659 | 732.11 | 705.8 | 68.15 | 0.54 | 25.93306 |
| 20.001 | 99 | 99 | ILSIIDKFEDNDPALHNVY | 739.3802 | 739.3793 | 3 | 87.39715 | 81762.3 | 77429.76 | 87.35 | 0.44 | 1530.255 |
| 21.001 | 99 | 99 | ILSIIDKFEDNDPALH | 613.9865 | 613.9877 | 3 | 81.68097 | 74473.22 | 73586.89 | 81.67 | 0.4 | 1055.993 |
| 22.001 | 99 | 99 | ILSIIDKFEDND | 711.3586 | 711.3616 | 2 | 79.60927 | 13195.17 | 13191.3 | 79.62 | 0.45 | 206.1864 |
| 24.001 | 99 | 99 | SIIDKFEDNDPALHNVY | 663.9894 | 663.9899 | 3 | 77.23023 | 47977.46 | 47089.07 | 77.31 | 0.5 | 405.0375 |

| | | | | | | | | | | | | |
|---|---|---|---|---|---|---|---|---|---|---|---|---|
| 25.001 | 99 | 99 | SIIDKFEDNDPALH | 538.5974 | 538.5984 | 3 | 67.05965 | 35644.68 | 30847.26 | 66.98 | 0.46 | 354.6286 |
| 26.001 | 99 | 99 | SIIDKFEDNDPALH | 807.8939 | 807.8859 | 2 | 67.00265 | 10029.18 | 9635.57 | 66.98 | 0.56 | 230.1213 |
| 27.001 | 99 | 99 | SIIDKFEDND | 598.2744 | 598.2775 | 2 | 59.29842 | 6678.93 | 5476.51 | 59.35 | 0.3 | 359.8172 |
| 29.001 | 24.8 | 24.83 | IDKFEDNDPALH | 707.3326 | 707.3359 | 2 | 67.10582 | 3245.46 | 3046.77 | 67.05 | 0.66 | 101.9649 |
| 31.001 | 99 | 99 | VFDIAPSVNASEIALF | 846.9433 | 846.9458 | 2 | 106.1601 | 79806.37 | 70157.22 | 106.2 | 0.16 | 1255.343 |
| 32.001 | 24.8 | 9.903 | VFDIAPSVN | 481.2491 | 481.2531 | 2 | 65.09825 | 3637.96 | 3447.75 | 65.09 | 0.63 | 38.33619 |
| 43.001 | 99 | 99 | IIGENYDAL | 504.2516 | 504.2558 | 2 | 64.73595 | 6474.71 | 2768.2 | 65.16 | 0.69 | 47.1384 |
| 44.001 | 99 | 99 | SYSDRESQSETVNY | 832.852 | 832.8553 | 2 | 40.43518 | 921.49 | 479.52 | 40.59 | 0.28 | 605.6223 |
| **45.001** | **99** | **99** | **SDRESQSETVNY** | **707.804** | **707.8077** | **2** | **32.77195** | **1357.66** | **1136.74** | **32.85** | **0.37** | **1792.464** |
| 46.001 | 99 | 99 | SDRESQSETVNY | 721.8197 | 721.8051 | 2 | 42.00142 | 405.32 | 399.31 | 41.98 | 0.38 | 982.1855 |
| 47.001 | 99 | 99 | SDRESQSETVNY | 708.2934 | 708.2997 | 2 | 34.80267 | 365.55 | 360.81 | 34.81 | 0.47 | 706.7414 |
| 48.001 | 99 | 99 | SDRESQSETVNY | 736.3109 | 736.3184 | 2 | 33.30592 | 301.94 | 248.1 | 33.25 | 0.4 | 583.4597 |
| 49.001 | 99 | 99 | EVSKGEELFTGVVPILVELDGDVNGHKF | 757.892 | 757.8933 | 4 | 106.0259 | 62228.4 | 46664.05 | 105.97 | 0.1 | 672.496 |
| 50.001 | 99 | 99 | EVSKGEELF | 519.2565 | 519.2611 | 2 | 52.8163 | 2292.46 | 1852.32 | 52.74 | 0.47 | 153.3652 |
| 51.001 | 54.3 | 54.27 | EVSKGEELF | 510.2896 | 510.2558 | 2 | 106.5314 | 549.75 | 546.89 | 106.52 | 0.17 | 30.33417 |
| 52.001 | 99 | 99 | FTGVVPILVELDGDVNGHKF | 719.7091 | 719.7124 | 3 | 104.2452 | 4916.05 | 4477.09 | 104.2 | 0.38 | 71.34451 |
| 53.001 | 99 | 99 | TGVVPILVELDGDVN | 770.4117 | 770.4169 | 2 | 102.0698 | 18070.43 | 17827.14 | 102.11 | 0.6 | 458.049 |
| 54.001 | 99 | 99 | TGVVPILVELDGDVNGHKF | 676.0158 | 676.0212 | 3 | 102.4589 | 5852.04 | 4608.9 | 102.72 | 0.71 | 116.5559 |
| 55.001 | 99 | 99 | TGVVPILVEL | 520.3186 | 520.3235 | 2 | 102.3253 | 3156.29 | 2544.56 | 102.24 | 0.43 | 59.15287 |
| 55.002 | 99 | 99 | VQERTIFF | 520.2796 | 520.2822 | 2 | 67.2188 | 3906.1 | 2121.15 | 67.41 | 0.5 | 57.14093 |
| 56.001 | 99 | 99 | TGVVPILVELDGDVNG | 798.9235 | 798.9276 | 2 | 102.4355 | 2389.77 | 2223.36 | 102.58 | 0.8 | 146.4749 |
| 57.001 | 99 | 99 | TGVVPILVELDGDVNGHKF | 670.6846 | 670.6896 | 3 | 115.4682 | 315.48 | 296.63 | 115.48 | 0.13 | 362.6304 |
| 65.001 | 99 | 99 | KFSVSGEGEGDATY | 723.819 | 723.8228 | 2 | 48.26007 | 1138.51 | 1040.44 | 48.4 | 0.44 | 351.4518 |
| 67.001 | 99 | 99 | SVSGEGEGDATY | 586.2377 | 586.2411 | 2 | 34.93982 | 7007.37 | 6997.59 | 34.93 | 0.45 | 7014.495 |
| 68.001 | 99 | 99 | SVSGEGEGDATY | 605.2109 | 605.2191 | 2 | 34.97782 | 3005.3 | 2914.57 | 34.93 | 0.45 | 3259.085 |
| 68.002 | 99 | 99 | SVSGEGEGDATY | 605.2109 | 605.2191 | 2 | 34.97782 | 3005.3 | 2914.57 | 34.93 | 0.45 | 3259.085 |
| 68.003 | 99 | 99 | SVSGEGEGDATY | 605.2109 | 605.2191 | 2 | 34.97782 | 3005.3 | 2914.57 | 34.93 | 0.45 | 3259.085 |
| 69.001 | 99 | 99 | SVSGEGEGDATY | 614.7454 | 614.7518 | 2 | 35.71562 | 1231.29 | 1142.17 | 35.76 | 0.71 | 1397.866 |
| 69.002 | 99 | 99 | SVSGEGEGDATY | 614.7454 | 614.7518 | 2 | 35.71562 | 1231.29 | 1142.17 | 35.76 | 0.71 | 1397.866 |
| 69.003 | 99 | 99 | SVSGEGEGDATY | 614.7454 | 614.7518 | 2 | 35.71562 | 1231.29 | 1142.17 | 35.76 | 0.71 | 1397.866 |
| 69.004 | 99 | 99 | SVSGEGEGDATY | 614.7454 | 614.7518 | 2 | 35.53945 | 1231.29 | 802.31 | 35.76 | 0.71 | 1288.663 |
| 69.005 | 99 | 25.83 | SVSGEGEGDATY | 614.7454 | 614.7518 | 2 | 35.53945 | 1231.29 | 802.31 | 35.76 | 0.71 | 1288.663 |

| | | | | | | | | | | | | |
|---|---|---|---|---|---|---|---|---|---|---|---|---|
| 69.006 | 99 | 25.83 | SVSGEGEGDATY | 614.7454 | 614.7518 | 2 | 35.53945 | 1231.29 | 802.31 | 35.76 | 0.71 | 1288.663 |
| 70.001 | 25.8 | 25.83 | SVSGEGEGDATY | 633.7216 | 633.7298 | 2 | 35.67795 | 505.75 | 461.58 | 35.72 | 0.71 | 685.1107 |
| 72.001 | 99 | 99 | ICTTGKLPVPW | 636.3392 | 636.3445 | 2 | 75.88865 | 3419.04 | 3002.65 | 75.94 | 0.49 | 87.66638 |
| 73.001 | 98.8 | 98.76 | ICTTGKLPVPWPTLVTTL | 999.0576 | 999.0606 | 2 | 102.1293 | 5936.46 | 5905.92 | 102.11 | 0.73 | 255.8852 |
| 77.001 | 99 | 99 | KSAMPEGYVQERTIF | 591.291 | 591.2943 | 3 | 57.84185 | 1426.92 | 1214.15 | 57.92 | 0.16 | 608.4799 |
| 78.001 | 99 | 99 | VQERTIF | 446.746 | 446.748 | 2 | 48.21505 | 3090.34 | 3085.36 | 48.2 | 0.37 | 499.5639 |
| 80.001 | 99 | 99 | KTRAEVKFEGDTL | 498.6037 | 498.6035 | 3 | 43.42933 | 67031.34 | 66730.55 | 43.42 | 0.37 | 44822.64 |
| 81.001 | 99 | 99 | KTRAEVKFEGDTLVN | 569.6402 | 569.6406 | 3 | 45.97402 | 19855.18 | 10504.87 | 45.8 | 0.33 | 2999.983 |
| 81.002 | 99 | 33.05 | KTRAEVKFEGDTLVNR | 569.6402 | 569.6406 | 3 | 45.65603 | 19855.18 | 10193.19 | 45.8 | 0.33 | 4222.191 |
| 82.001 | 99 | 99 | KTRAEVKFEGDTL | 517.6104 | 517.6106 | 3 | 43.8683 | 10828.41 | 10402.23 | 43.92 | 0.38 | 8923.275 |
| 82.002 | 99 | 99 | KTRAEVKFEGDTL | 517.6104 | 517.6106 | 3 | 43.8683 | 10828.41 | 10402.23 | 43.92 | 0.38 | 8923.275 |
| 82.003 | 99 | 99 | KTRAEVKFEGDTL | 517.6104 | 517.6106 | 3 | 43.61598 | 10828.41 | 4577.42 | 43.92 | 0.38 | 4628.193 |
| 82.004 | 99 | 99 | KTRAEVKFEGDTL | 517.6104 | 517.6106 | 3 | 43.61598 | 10828.41 | 4577.42 | 43.92 | 0.38 | 4628.193 |
| 82.005 | 99 | 99 | KTRAEVKFEGDTL | 517.6104 | 517.6106 | 3 | 43.61598 | 10828.41 | 4577.42 | 43.92 | 0.38 | 4628.193 |
| 82.006 | 99 | 99 | KTRAEVKFEGDTL | 517.6104 | 517.6106 | 3 | 43.61598 | 10828.41 | 4577.42 | 43.92 | 0.38 | 4628.193 |
| 82.007 | 99 | 99 | KTRAEVKFEGDTL | 517.6091 | 517.6106 | 3 | 44.6751 | 4767.43 | 4727.15 | 44.69 | 0.42 | 2190.815 |
| 82.008 | 99 | 25.83 | KTRAEVKFEGDTL | 517.6104 | 517.6106 | 3 | 43.8683 | 10828.41 | 10402.23 | 43.92 | 0.38 | 8923.275 |
| 83.001 | 99 | 99 | KTRAEVKFEGDTLVNRIEL | 740.0755 | 740.0779 | 3 | 64.97377 | 8857.75 | 8580.48 | 65.02 | 0.36 | 223.7829 |
| 84.001 | 99 | 99 | KTRAEVKFEGDTLVN | 588.6455 | 588.6477 | 3 | 46.00868 | 3112.78 | 2892.93 | 46.16 | 0.69 | 1507.25 |
| 84.002 | 99 | 99 | KTRAEVKFEGDTLVN | 588.6455 | 588.6477 | 3 | 46.00868 | 3112.78 | 2892.93 | 46.16 | 0.69 | 1507.25 |
| 84.003 | 99 | 99 | KTRAEVKFEGDTLVN | 588.6455 | 588.6477 | 3 | 46.00868 | 3112.78 | 2892.93 | 46.16 | 0.69 | 1507.25 |
| 84.004 | 99 | 99 | KTRAEVKFEGDTLVN | 588.6455 | 588.6477 | 3 | 46.00868 | 3112.78 | 2892.93 | 46.16 | 0.69 | 1507.25 |
| 84.005 | 99 | 99 | KTRAEVKFEGDTLVN | 588.6455 | 588.6477 | 3 | 46.00868 | 3112.78 | 2892.93 | 46.16 | 0.69 | 1507.25 |
| 84.006 | 99 | 99 | KTRAEVKFEGDTLVN | 588.6455 | 588.6477 | 3 | 46.00868 | 3112.78 | 2892.93 | 46.16 | 0.69 | 1507.25 |
| 84.007 | 99 | 99 | KTRAEVKFEGDTLVN | 588.6455 | 588.6477 | 3 | 46.00868 | 3112.78 | 2892.93 | 46.16 | 0.69 | 1507.25 |
| 84.008 | 99 | 99 | KTRAEVKFEGDTLVN | 588.6455 | 588.6477 | 3 | 46.00868 | 3112.78 | 2892.93 | 46.16 | 0.69 | 1507.25 |
| 85.001 | 99 | 99 | KTRAEVKFEGDTLV | 787.9294 | 787.9305 | 2 | 46.05018 | 645.69 | 367.5 | 45.84 | 0.46 | 918.7012 |
| 85.002 | 99 | 99 | KTRAEVKFEGDTLV | 787.9294 | 787.9305 | 2 | 46.05018 | 645.69 | 367.5 | 45.84 | 0.46 | 918.7012 |
| 86.001 | 23.9 | 23.9 | KTRAEVKFEGD | 631.3271 | 631.3304 | 2 | 43.34668 | 1323.54 | 1177.54 | 43.42 | 0.47 | 1375.873 |
| 86.002 | 23.9 | 23.38 | KTRAEVKFEGD | 631.3271 | 631.3304 | 2 | 43.34668 | 1323.54 | 1177.54 | 43.42 | 0.47 | 1375.873 |
| 87.001 | 23.4 | 23.38 | KTRAEVKFEGDTL | 503.2605 | 503.2595 | 3 | 43.49434 | 5561.58 | 4569.87 | 43.42 | 0.41 | 3046.753 |
| 88.001 | 23.4 | 23.38 | KTRAEVKFEGDTL | 497.9335 | 497.9316 | 3 | 43.92663 | 2569.09 | 2523.18 | 43.92 | 0.38 | 3460.399 |

| | | | | | | | | | | | | |
|---|---|---|---|---|---|---|---|---|---|---|---|---|
| 90.001 | 99 | 99 | TRAEVKFEGDTL | 455.9035 | 455.9052 | 3 | 50.09478 | 862.41 | 850.39 | 50.08 | 0.3 | 272.6838 |
| 92.001 | 99 | 99 | AEVKFEGDTL | 554.7775 | 554.7797 | 2 | 56.41325 | 8159.77 | 7906.75 | 56.43 | 0.3 | 553.6661 |
| 93.001 | 24.8 | 24.81 | EVKFEGDTL | 510.2574 | 510.2558 | 2 | 73.1568 | 20185.66 | 16328.21 | 73.28 | 0.4 | 199.3768 |
| 94.001 | 99 | 99 | EGDTLVNRIEL | 629.8312 | 629.8355 | 2 | 70.72877 | 21060.21 | 20491.92 | 70.75 | 0.27 | 240.7872 |
| 96.001 | 99 | 99 | VNRIEL | 372.2207 | 372.2242 | 2 | 46.83247 | 977 | 647.72 | 46.92 | 0.33 | 71.18753 |
| 97.001 | 99 | 99 | KGIDFKEDGNIL | 450.2393 | 450.2418 | 3 | 61.50563 | 69062.35 | 68286.92 | 61.5 | 0.23 | 1860.696 |
| 98.001 | 99 | 99 | KGIDFKEDGNILGH | 514.9346 | 514.9352 | 3 | 55.48032 | 44621.95 | 40695.55 | 55.53 | 0.33 | 1311.657 |
| 102.001 | 28.8 | 28.83 | EYNYNSHNVY | 652.2744 | 652.265 | 2 | 38.8191 | 418.67 | 341.01 | 38.94 | 0.44 | 528.7459 |
| 102.002 | 28.8 | 28.83 | EYNYNSHNVY | 652.2744 | 652.265 | 2 | 38.8191 | 418.67 | 341.01 | 38.94 | 0.44 | 528.7459 |
| 102.003 | 28.8 | 28.83 | EYNYNSHNVY | 652.2744 | 652.265 | 2 | 38.8191 | 418.67 | 341.01 | 38.94 | 0.44 | 528.7459 |
| 105.001 | 99 | 99 | KIRHNIEDGSVQLADHY | 499.5074 | 499.5065 | 4 | 43.95963 | 31025.07 | 30000.06 | 43.98 | 0.44 | 10583.49 |
| 105.002 | 99 | 99 | IRHNIEDGSVQLADHY | 399.805 | 399.8067 | 5 | 43.9873 | 2697.63 | 2512.53 | 43.98 | 0.38 | 1911.108 |
| 105.003 | 99 | 23.38 | KIRHNIEDGSVQLADHY | 665.6741 | 665.6729 | 3 | 43.93997 | 25258.4 | 23039.5 | 43.98 | 0.41 | 17238.88 |
| 106.001 | 99 | 99 | KIRHNIEDGSVQLADHY | 513.7635 | 513.7619 | 4 | 43.83498 | 10282.79 | 10236.92 | 43.85 | 0.25 | 9867.714 |
| 106.002 | 99 | 99 | KIRHNIEDGSVQLADHY | 513.7605 | 513.7619 | 4 | 43.30684 | 2474.96 | 2410.48 | 43.35 | 0.28 | 5369.28 |
| 106.003 | 99 | 99 | KIRHNIEDGSVQLADHY | 513.7605 | 513.7619 | 4 | 43.30684 | 2474.96 | 2410.48 | 43.35 | 0.28 | 5369.28 |
| 106.004 | 99 | 99 | KIRHNIEDGSVQLADHY | 513.7635 | 513.7619 | 4 | 43.76982 | 10282.79 | 8293.24 | 43.85 | 0.25 | 7516.121 |
| 106.005 | 99 | 99 | KIRHNIEDGSVQLADHY | 513.7635 | 513.7619 | 4 | 43.83498 | 10282.79 | 10236.92 | 43.85 | 0.25 | 9867.714 |
| 106.006 | 99 | 99 | KIRHNIEDGSVQLADHY | 513.7635 | 513.7619 | 4 | 43.83498 | 10282.79 | 10236.92 | 43.85 | 0.25 | 9867.714 |
| 106.007 | 99 | 99 | KIRHNIEDGSVQLADHY | 513.7635 | 513.7619 | 4 | 43.76982 | 10282.79 | 8293.24 | 43.85 | 0.25 | 7516.121 |
| 106.008 | 99 | 99 | KIRHNIEDGSVQLADHY | 513.7635 | 513.7619 | 4 | 43.76982 | 10282.79 | 8293.24 | 43.85 | 0.25 | 7516.121 |
| 106.009 | 99 | 99 | KIRHNIEDGSVQLADHY | 513.7605 | 513.7619 | 4 | 43.30684 | 2474.96 | 2410.48 | 43.35 | 0.28 | 5369.28 |
| 107.001 | 99 | 99 | KIRHNIEDGSVQLADHY | 503.2543 | 503.2592 | 4 | 43.9613 | 7048.29 | 6896.94 | 43.98 | 0.41 | 5954.47 |
| 108.001 | 99 | 99 | KIRHNIEDGSVQLADHY | 671.002 | 671.0046 | 3 | 43.94164 | 5716.65 | 5498.52 | 43.98 | 0.19 | 3435.005 |
| 108.002 | 99 | 99 | KIRHNIEDGSVQLADHY | 671.002 | 671.0046 | 3 | 43.94164 | 5716.65 | 5498.52 | 43.98 | 0.19 | 3435.005 |
| 109.001 | 99 | 99 | KIRHNIEDGSVQLADHY | 499.754 | 499.7525 | 4 | 45.7852 | 5135.04 | 4955.75 | 45.8 | 0.23 | 1965.622 |
| 109.002 | 99 | 99 | KIRHNIEDGSVQLADHY | 665.9996 | 666.0009 | 3 | 45.75787 | 3569.48 | 3120.9 | 45.8 | 0.26 | 1680.25 |
| 110.001 | 99 | 99 | KIRHNIEDGSVQLADHY | 514.0128 | 514.0079 | 4 | 43.9663 | 5330.03 | 3743.01 | 43.89 | 0.47 | 5443.799 |
| 110.002 | 99 | 99 | KIRHNIEDGSVQLADHY | 514.0128 | 514.0079 | 4 | 43.9663 | 5330.03 | 3743.01 | 43.89 | 0.47 | 5443.799 |
| 110.003 | 99 | 99 | KIRHNIEDGSVQLADHY | 514.0128 | 514.0079 | 4 | 43.9663 | 5330.03 | 3743.01 | 43.89 | 0.47 | 5443.799 |
| 110.004 | 99 | 33.05 | KIRHNIEDGSVQLADHY | 514.0128 | 514.0079 | 4 | 43.9663 | 5330.03 | 3743.01 | 43.89 | 0.47 | 5443.799 |
| 110.005 | 99 | 27.51 | KIRHNIEDGSVQLADHY | 514.0128 | 514.0079 | 4 | 43.9663 | 5330.03 | 3743.01 | 43.89 | 0.47 | 5443.799 |

| | | | | | | | | | | | | |
|---|---|---|---|---|---|---|---|---|---|---|---|---|
| 110.006 | 99 | 23.38 | KIRHNIEDGSVQLADHY | 514.0128 | 514.0079 | 4 | 43.9663 | 5330.03 | 3743.01 | 43.89 | 0.47 | 5443.799 |
| 111.001 | 99 | 97.36 | KIRHNIEDGSVQLADHY | 671.0049 | 671.0046 | 3 | 50.61125 | 2726.36 | 2662.74 | 50.62 | 0.31 | 310.7416 |
| 112.001 | 99 | 99 | KIRHNIEDGS | 575.7991 | 575.8018 | 2 | 44.02363 | 2378.62 | 2302.35 | 44.01 | 0.44 | 2335.815 |
| 113.001 | 99 | 99 | KIRHNIEDGSVQLADHY | 678.3201 | 678.3249 | 3 | 44.00397 | 1587.2 | 1471.72 | 43.98 | 0.22 | 1929.413 |
| 113.002 | 99 | 99 | KIRHNIEDGSVQLADHY | 678.3201 | 678.3249 | 3 | 44.00397 | 1587.2 | 1471.72 | 43.98 | 0.22 | 1929.413 |
| 114.001 | 99 | 99 | KIRHNIEDGSVQLADHY | 528.0153 | 528.0173 | 4 | 43.14118 | 1419.98 | 1406.58 | 43.13 | 0.42 | 2279.413 |
| 115.001 | 99 | 99 | KIRHNIEDGSVQLADHY | 675.3229 | 675.3447 | 3 | 43.97298 | 1367.03 | 1364.88 | 43.98 | 0.5 | 2150.469 |
| 115.002 | 99 | 99 | KIRHNIEDGSVQLADHY | 675.3229 | 675.3447 | 3 | 43.97298 | 1367.03 | 1364.88 | 43.98 | 0.5 | 2150.469 |
| 115.003 | 99 | 99 | KIRHNIEDGSVQLADHY | 675.3229 | 675.3326 | 3 | 43.97298 | 1367.03 | 1364.88 | 43.98 | 0.5 | 2150.469 |
| 115.004 | 99 | 99 | KIRHNIEDGSVQLADHY | 675.3229 | 675.3326 | 3 | 43.97298 | 1367.03 | 1364.88 | 43.98 | 0.5 | 2150.469 |
| 116.001 | 99 | 99 | KIRHNIEDGSVQLADHY | 703.6824 | 703.6873 | 3 | 43.15619 | 750.89 | 713.21 | 43.13 | 0.51 | 1266.144 |
| 116.002 | 99 | 99 | KIRHNIEDGSVQLADHY | 703.6824 | 703.6873 | 3 | 43.15619 | 750.89 | 713.21 | 43.13 | 0.51 | 1266.144 |
| 116.003 | 99 | 99 | KIRHNIEDGSVQLADHY | 703.6824 | 703.6873 | 3 | 43.15619 | 750.89 | 713.21 | 43.13 | 0.51 | 1266.144 |
| 116.004 | 99 | 99 | KIRHNIEDGSVQLADHY | 703.6824 | 703.6873 | 3 | 43.15619 | 750.89 | 713.21 | 43.13 | 0.51 | 1266.144 |
| 116.005 | 99 | 99 | KIRHNIEDGSVQLADHY | 703.6824 | 703.6873 | 3 | 43.15619 | 750.89 | 713.21 | 43.13 | 0.51 | 1266.144 |
| 116.006 | 99 | 99 | KIRHNIEDGSVQLADHY | 703.6824 | 703.6873 | 3 | 43.15619 | 750.89 | 713.21 | 43.13 | 0.51 | 1266.144 |
| 116.007 | 99 | 99 | KIRHNIEDGSVQLADHY | 703.6824 | 703.6873 | 3 | 43.15619 | 750.89 | 713.21 | 43.13 | 0.51 | 1266.144 |
| 116.008 | 99 | 99 | KIRHNIEDGSVQLADHY | 703.6824 | 703.6873 | 3 | 43.15619 | 750.89 | 713.21 | 43.13 | 0.51 | 1266.144 |
| 116.009 | 99 | 27.51 | KIRHNIEDGSVQLADHY | 703.6824 | 703.6873 | 3 | 43.15619 | 750.89 | 713.21 | 43.13 | 0.51 | 1266.144 |
| 116.010 | 99 | 25.83 | KIRHNIEDGSVQLADHY | 703.6824 | 703.6873 | 3 | 43.15619 | 750.89 | 713.21 | 43.13 | 0.51 | 1266.144 |
| 116.011 | 99 | 25.83 | KIRHNIEDGSVQLADHY | 703.6824 | 703.6873 | 3 | 43.15619 | 750.89 | 713.21 | 43.13 | 0.51 | 1266.144 |
| 116.012 | 99 | 23.38 | KIRHNIEDGSVQLADHY | 703.6824 | 703.6873 | 3 | 43.15619 | 750.89 | 713.21 | 43.13 | 0.51 | 1266.144 |
| 116.013 | 99 | 23.38 | KIRHNIEDGSVQLADHY | 703.6824 | 703.6873 | 3 | 43.15619 | 750.89 | 713.21 | 43.13 | 0.51 | 1266.144 |
| 117.001 | 49.2 | 24.85 | KIRHNIEDGSVQL | 503.933 | 503.9388 | 3 | 41.7246 | 381.94 | 343.22 | 41.69 | 0.42 | 1012.293 |
| 118.001 | 30.9 | 30.91 | KIRHNIEDGSVQLADHYQ | 702.3154 | 702.3556 | 3 | 43.83998 | 1571.33 | 1491.09 | 43.92 | 0.64 | 2511.26 |
| 119.001 | 28.5 | 28.51 | KIRHNIEDGSV | 625.3333 | 625.336 | 2 | 43.97132 | 2426.35 | 2376.31 | 43.98 | 0.44 | 1914.609 |
| 120.001 | 23.4 | 23.38 | KIRHNIEDGSVQLADHY | 683.3094 | 683.3447 | 3 | 44.00563 | 4894.15 | 4738.7 | 43.98 | 0.44 | 5581.833 |
| 123.001 | 99 | 99 | HNIEDGSVQLADHY | 533.2446 | 533.2462 | 3 | 52.01235 | 1927.2 | 1862.88 | 51.96 | 0.34 | 236.48 |
| 124.001 | 98.6 | 98.56 | HNIEDGSVQLADHY | 799.8668 | 799.8577 | 2 | 51.95285 | 692.2 | 682.47 | 51.96 | 0.47 | 116.2673 |
| 125.001 | 99 | 98.56 | NIEDGSVQLADHY | 730.8357 | 730.8362 | 2 | 59.0546 | 1934.51 | 1886.96 | 59.08 | 0.65 | 227.5839 |
| 126.001 | 37.6 | 37.61 | EDGSVQLADHY | 617.2676 | 617.2728 | 2 | 44.02697 | 307.19 | 289.62 | 44.01 | 0.28 | 1084.4 |
| 127.001 | 17.7 | 17.67 | EDGSVQLADHY | 608.3057 | 608.2675 | 2 | 44.15396 | 183.75 | 143.36 | 43.82 | 0.44 | 915.3699 |

| | | | | | | | | | | | | |
|---|---|---|---|---|---|---|---|---|---|---|---|---|
| 130.001 | 99 | 99 | QQNTPIGDGPVLLPDNHY | 980.9722 | 980.9736 | 2 | 80.15307 | 28063.26 | 22907.35 | 80.24 | 0.41 | 415.8313 |
| 131.001 | 99 | 99 | QQNTPIGDGPVLLPDNHY | 989.4828 | 989.4869 | 2 | 70.50661 | 23628.15 | 22334.36 | 70.58 | 0.37 | 327.209 |
| 134.001 | 99 | 99 | EFVTAAGITL | 511.2795 | 511.2819 | 2 | 84.47082 | 74461.3 | 74078.02 | 84.43 | 0.33 | 640.788 |
| 135.001 | 99 | 99 | EFVTAAGITL | 502.2599 | 502.2766 | 2 | 73.1288 | 30227.32 | 19371.15 | 73.24 | 0.4 | 264.5301 |
| 135.002 | 99 | 24.8 | LEFVTAAGIT | 502.2599 | 502.2766 | 2 | 73.1288 | 30227.32 | 19371.15 | 73.24 | 0.4 | 264.5301 |
| 136.001 | 99 | 99 | EFVTAAGITLGMDELYK | 937.4659 | 937.4662 | 2 | 81.78747 | 32609.8 | 18645.39 | 82 | 0.51 | 488.0495 |
| 137.001 | 99 | 99 | EFVTAAGITLGMDELYK | 929.4674 | 929.4688 | 2 | 71.80372 | 4975.33 | 4843.17 | 71.75 | 0.41 | 180.572 |
| 139.001 | 99 | 99 | VTAAGITLGMDELYK | 799.4073 | 799.4107 | 2 | 67.96777 | 9607.54 | 8896.47 | 68.05 | 0.44 | 194.3755 |
| 141.001 | 99 | 99 | FCQVGYTL | 494.2625 | 494.2339 | 2 | 73.12714 | 60588.73 | 35799.52 | 73.24 | 0.4 | 423.253 |
| 142.001 | 99 | 99 | LAIESANVSPAI | 592.8441 | 592.8297 | 2 | 79.63343 | 7496.75 | 6752.05 | 79.69 | 0.31 | 251.8267 |
| 143.001 | 99 | 99 | DIGKLEIRNVL | 442.928 | 442.9295 | 3 | 47.88258 | 2908.19 | 2465.86 | 47.97 | 0.37 | 424.0702 |
| 144.001 | 99 | 99 | QVSGDEINHRI | 634.3336 | 634.3231 | 2 | 61.89812 | 2674.41 | 2262.25 | 61.85 | 0.63 | 85.36069 |
| 145.001 | 98 | 98.01 | DKQDQNLENSLSL | 771.3585 | 771.3458 | 2 | 114.6649 | 301.49 | 295.08 | 114.68 | 0.16 | 262.0085 |
| 146.001 | 93.4 | 93.38 | KVEARTKY | 497.7615 | 497.7876 | 2 | 49.23034 | 1394.76 | 990.49 | 49.34 | 0.44 | 228.9957 |
| 147.001 | 26.6 | 26.62 | NGDDKFF | 441.2138 | 441.1931 | 2 | 45.65103 | 890.9 | 513.7 | 45.84 | 0.69 | 1097.603 |
| 148.001 | 25.1 | 25.12 | DKSLKSQT | 453.7238 | 453.7482 | 2 | 40.83265 | 416.71 | 402.28 | 40.84 | 0.59 | 1620.935 |
| 149.001 | 24.8 | 24.82 | SFKHGNVDGDLEVLIP | 870.4472 | 870.4518 | 2 | 66.33485 | 60463.32 | 44824.98 | 66.45 | 0.4 | 685.7784 |
| 150.001 | 24.8 | 24.81 | KNLIVIESQSETVNYE | 622.6545 | 622.6545 | 3 | 67.35547 | 7023.81 | 5957.83 | 67.41 | 0.36 | 161.1039 |
| 151.001 | 21.1 | 21.14 | INGDEKF | 411.7186 | 411.7032 | 2 | 41.07413 | 1383.31 | 886.4 | 40.84 | 0.53 | 5638.984 |
| 152.001 | 17.7 | 17.67 | KNLIVIESQSETVNY | 435.4801 | 435.4741 | 4 | 45.7462 | 769.22 | 741.55 | 45.77 | 0.33 | 921.9589 |
